# Prefrontal cortex transcriptomes distinguish Alzheimer’s disease from asymptomatic amyloid pathology, neuropeptide loss, immune-complement activation and glial remodelling

**DOI:** 10.64898/2026.10.31.763276

**Authors:** Yusuf Munir Aliyu, Umar Sagir, Fahad Sajjad, Sania Khurshid, Farha Tarique, Ibrahim Sani Maikudi

**Affiliations:** Department of General Studies, Kasimu Kofar Bai, College of Nursing Sciences, Katsina, Nigeria; Department of Microbiology, Umaru Musa Yar’adua University, PMB 2218, Katsina, Nigeria; Department of Biological Sciences, Federal University Dutsin-Ma, Nigeria; Institute of Biotechnology and Genetic Engineering, Agriculture University of Peshawar, Peshawar, Pakistan; Department of Medical Laboratory Technology, National University of Medical Sciences, Rawalpindi, Pakistan; Centre of Bioinformatics, Institute of Interdisciplinary Studies, University of Allahabad, Allahabad, India; Department of Biotechnology, Federal University of Medical and Health Sciences, Funtua, Nigeria

**Keywords:** Dorsolateral prefrontal cortex, RNA-seq, differential expression, neuropeptide signalling, myelination, *APOE*, *TREM2*

## Abstract

Alzheimer’s disease (AD) is now defined biologically by cortical amyloid-β (A) and tau (T) pathology, yet the transcriptional programmes that separate symptomatic AD (A+T+) from amyloid-positive, tau-negative pathology ageing (A+T−) and from pathology-free controls (A−T−) remain incompletely resolved in the association cortex. We re-analysed bulk RNA-sequencing data from the dorsolateral prefrontal cortex (DLPFC) of 15 donors (BioProject PRJNA1017546; 87 sequencing runs aggregated to donor level): Normal (n = 2), pathology ageing (n = 5) and AD (n = 8). Differential expression was tested with DESeq2 (adjusted P < 0.05, |log fold change| > 1), and up- and down-regulated gene sets were analysed separately by Gene Ontology (GO) and KEGG over-representation. APOE and TREM2 p.Arg47His contrasts were treated as exploratory. AD versus Normal yielded 341 differentially expressed genes (DEGs; 223 up, 118 down) and AD versus pathology ageing yielded 496 (358 up, 138 down), whereas pathology ageing versus Normal yielded only six. Genes reduced in AD were dominated by neuropeptide and hormone signalling (*VGF*, *VIP*, *CRH*, *GRP*, *TAC1*, *PENK*; neuroactive ligand–receptor interaction, adjusted P = 8.8 × 10; neuropeptide hormone activity, adjusted P = 6.3 × 10), most of which also differed from pathology ageing. Genes increased in AD were enriched for interferon-γ response, myeloid activation, adaptive immunity and classical complement (*C1QA*, *C1QB*, *C1QC*). Relative to pathology ageing, AD additionally showed induction of gliogenesis, oligodendrocyte differentiation and myelination programmes (gliogenesis, 21 genes, adjusted P = 2.6 × 10; *SOX10*, *MYRF*) and of the microglial purinergic receptor gene *P2RX7* (log FC = 1.46, adjusted P = 1.5 × 10). All six pathology-ageing DEGs, including the myeloid genes *VSIG4* and *CD163*, were also DEGs in AD, and 50 DEGs shared by the two AD contrasts moved in the same direction. Genotype-stratified analyses were underpowered or confounded with diagnosis. In the DLPFC, AD is therefore characterised by a graded loss of neuropeptidergic signalling, myeloid–complement activation that begins at the pathology-ageing stage, and an AD-specific glial/oligodendrocyte response. These signatures nominate candidate biomarkers and mechanisms for early intervention, but the small cohort demands replication in larger, covariate-annotated datasets.

## 1.0 Introduction

Alzheimer’s disease is the leading cause of dementia worldwide and is characterised neuropathologically by extracellular amyloid-β plaques and intraneuronal neurofibrillary tangles of hyperphosphorylated tau (Machlovi *et al*., 2022; Brase *et al*., 2023). The 2018 NIA-AA Research Framework recast AD as a biological continuum defined by the A/T/N biomarker scheme, so that individuals can be positioned along the disease trajectory by the presence of amyloid (A) and tau (T) pathology independently of their clinical status (Hampel *et al*., 2021). Within this framework, neuropathological and biomarker studies frequently identify cognitively unimpaired older individuals with abnormal amyloid but little or no neocortical tau, corresponding to an A+T− state often described as Alzheimer’s pathologic change, preclinical AD, or asymptomatic amyloidosis (Rafii *et al*., 2020; Leeuw *et al*., 2025). Longitudinal biomarker studies further support the view that amyloid positivity generally precedes substantial tau abnormality and cognitive decline, making the transition from amyloid-only states to tau-positive disease a central stage for mechanistic investigation (Leeuw *et al*., 2025).

Transcriptomic profiling of post-mortem brain has been a productive route to such mechanisms. Co-expression analyses of large human cortical datasets have implicated immune and microglial modules in late-onset AD and disease progression (Malamon & Kriete, 2020). Single-cell and single-nucleus studies have refined these signals to cell-type-specific states, including disease-associated microglia, complement-linked microglial programmes, oligodendroglial lysosomal dysfunction, and vascular or endothelial changes (Brase *et al*., 2023; Olayinka *et al*., 2025; Liu *et al*., 2026). Genome-wide association studies converge on the same biology: AD risk loci are enriched for microglial, immune, and protein catabolism/endolysosomal pathways, reinforcing innate immunity and lipid handling as causal contributors to disease risk (Wightman *et al*., 2021; Bellenguez *et al*., 2022; Brase *et al*., 2023). The APOE ε4 allele is the strongest common genetic risk factor for sporadic AD, and the TREM2 R47H variant is a major low-frequency risk allele linked to microglial dysfunction and disease-associated transcriptional responses (Machlovi *et al*., 2022; Brase *et al*., 2023; Carling *et al*., 2024). Despite this progress, key questions remain open for the association cortex, which is vulnerable to both amyloid and tau and supports higher-order cognitive functions. Separating transcriptional changes related specifically to the transition from amyloid-only pathology to full AD remains important because amyloid-positive asymptomatic individuals already show measurable risk for later tau accumulation and cognitive decline, but tau burden is more tightly linked to short-term clinical progression (Ossenkoppele *et al*., 2022; Gallego-Rudolf *et al*., 2024; Sperling *et al*., 2024). This also motivates testing whether neuronal programmes decline while glial and myeloid programmes increase across A−T−, A+T−, and A+T+ states, consistent with evidence that amyloid and tau interact with distinct but converging neurobiological processes across preclinical AD (Ossenkoppele *et al*., 2022). It is likewise reasonable to ask whether APOE ε4- and TREM2-linked effects remain visible in bulk prefrontal transcriptomes after accounting for pathology stage, given evidence that these variants alter cell-specific programmes involving ferroptosis, complement, autophagy-lysosomal pathways, interferon signalling, and microglial senescence (Machlovi *et al*., 2022; Brase *et al*., 2023; Carling *et al*., 2024).

Here we address these questions using bulk RNA-sequencing data from the dorsolateral prefrontal cortex (DLPFC) of donors stratified by A/T pathology. We quantify differential expression among Normal, pathology-ageing, and AD brains; characterise the biological processes and pathways represented by dysregulated genes; identify changes shared across disease contrasts; and explore genotype-stratified contrasts for APOE and TREM2. This framing fits current AD models in which amyloid-positive, tau-negative states can precede broader tau spread, neurodegeneration, immune activation, and symptomatic decline, making DLPFC transcriptomics a relevant approach for studying progression biology.

## 2.0 Materials and Methods

### 2.1 Study Cohort and Data Source

We analysed publicly available RNA-sequencing data deposited under BioProject PRJNA1017546 (SRA study SRP460672; University of Pennsylvania sequencing centre; released 2 October 2023). The dataset comprises Homo sapiens dorsolateral prefrontal cortex tissue sequenced on an Illumina platform with paired-end reads, a standard format also used in large human DLPFC transcriptomic studies (Prater *et al*., 2023; Green *et al*., 2024). In total, 87 sequencing runs representing 15 donors were available. Donor-level metadata comprised neuropathological diagnosis, A/T score, APOE genotype and TREM2 genotype (Table 1). Age, sex, post-mortem interval, RNA integrity and Braak/CERAD scores were not available in the files supplied for this analysis. The data are de-identified and publicly released; public-access brain transcriptomic resources commonly distribute de-identified data under privacy-protecting access frameworks for secondary analysis.

**Table 1.** Characteristics of the donor cohort by neuropathological group.

| Characteristic | Normal (A−T−) | Pathology ageing (A+T−) | Alzheimer’s disease (A+T+) | Total |
|---|---|---|---|---|
| Donors, *n* | 2 | 5 | 8 | 15 |
| *APOE* E3/E3, *n* | 2 | 1 | 2 | 5 |
| *APOE* E3/E4, *n* | 0 | 4 | 4 | 8 |
| *APOE* E4/E4, *n* | 0 | 0 | 2 | 2 |
| *TREM2* wild type, *n* | 2 | 4 | 5 | 11 |
| *TREM2* p.Arg47His carrier, *n* | 0 | 1 | 3 | 4 |
| Tissue | DLPFC | DLPFC | DLPFC | DLPFC |
| Platform/layout | HiSeq 2500, paired-end | HiSeq 2500, paired-end | HiSeq 2500, paired-end | 87 runs |

The 15 donors whose dorsolateral prefrontal cortex RNA-seq profiles were analysed, partitioned by diagnosis/A-T score, with the distribution of *APOE* and *TREM2* genotypes in each group. It documents the group sizes underlying every contrast and the degree to which genotype is confounded with diagnosis.A, amyloid-β pathology; T, tau pathology; DLPFC, dorsolateral prefrontal cortex. The 87 sequencing runs were summed per donor before analysis. Age, sex, post-mortem interval and RNA integrity were not available.

### 2.2 Group Definitions

Donors were assigned to three groups according to the diagnosis and A/T score provided with the dataset: Normal (A−T−; n = 2), pathology ageing (PA; A+T−; n = 5) and Alzheimer’s disease (AD; A+T+; n = 8). The APOE genotype was E3/E3 (n = 5), E3/E4 (n = 8) or E4/E4 (n = 2), and TREM2 was wild type (n = 11) or p.Arg47His (R47H) carrier (n = 4). No Normal donor carried an ε4 allele or the R47H variant, and both E4/E4 donors were AD cases (Table 1). These cohort-specific counts come from the deposited dataset and generally should not be literature-cited unless you also cite the dataset record itself.

#### 2.3.1 Read Quality Control, Mapping and Quantification

The data pre-processing was conducted in <u>UseGalaxy</u>. Read quality was assessed with FastQC (Galaxy Version 0.74+galaxy1) and summarised with MultiQC (Galaxy Version 1.35+galaxy4), and adapter and quality trimming were performed with fastp (Galaxy Version 1.4.0+galaxy0) and aligned using HISAT2 (Galaxy Version 2.2.3+galaxy0), Gene-level read counts for each run were generated with featureCounts, which is a general-purpose read summarisation program for assigning aligned reads to genomic features and supports both single-end and paired-end RNA-seq data (Liao *et al*., 2013). Gene-level summarisation by counting reads overlapping annotated exons for each gene is a common RNA-seq strategy, typically using RefSeq or Ensembl-style annotations (Liao *et al*., 2013). Read counting and read summarisation are essential steps in RNA-seq workflows, and differential expression analyses are often performed using total reads per gene (Schmid & Grossniklaus, 2015). Counts from runs belonging to the same donor were summed to obtain one donor-level count vector per donor; aggregation of counts across lower-level units into a sample-level profile is also used in transcriptomic workflows such as pseudobulk analysis (Green *et al*., 2024; Oxendine *et al*., 2025).

### 2.4 Exploratory Analysis

Principal component analysis (PCA) of DESeq2-normalised, variance-stabilised counts was used to examine global sample structure by diagnosis. PCA is a standard exploratory step in RNA-seq workflows and is commonly used to assess sample clustering and detect outlier structure in brain transcriptomic datasets (Olayinka *et al*., 2025; Vetrivel, 2026). DESeq2 provides transformed count data for sample-level exploratory analyses, and variance-stabilising or related transformations are used to reduce mean-dependent variance before clustering or ordination (Mathys *et al*., 2023). We inspected DESeq2 dispersion estimates to confirm the expected mean–dispersion trend, a standard diagnostic of negative-binomial count modelling (Prater *et al*., 2023).

### 2.5 Differential Expression Analysis

Differential expression was assessed with DESeq2, which model’s RNA-seq count data with negative-binomial generalized linear models (Mathys *et al*., 2023; Prater *et al*., 2023). Negative-binomial methods are widely used for RNA-seq differential expression because they explicitly model count structure and dispersion across genes (Schmid & Grossniklaus, 2015). DESeq2 was applied to test pairwise contrasts among diagnosis groups after pre-filtering low-count genes. P values were adjusted for multiple testing by the Benjamini–Hochberg procedure, which controls the false discovery rate in high-dimensional testing (Brandon *et al*., 2024; Green *et al*., 2024). DESeq2’s independent filtering is consistent with common RNA-seq practice in which lowly expressed genes are removed before or during multiple-testing adjustment to improve power (Ziemann *et al*., 2024; Olayinka *et al*., 2025). A gene was called differentially expressed when adjusted P < 0.05 and |log fold change| > 1, a common thresholding strategy in downstream DEG and enrichment workflows (Ziemann *et al*., 2024; Stead *et al*., 2025).

### 2.6 Exploratory Genotype Analyses

Secondary contrasts were run to explore genotype effects: APOE E4/E4 vs E3/E3, APOE E3/E4 vs E3/E3 and TREM2 R47H vs wild type. Because genotype is strongly correlated with diagnosis in this small cohort, these contrasts were not adjusted for diagnosis and are interpreted as hypothesis-generating only. Small-cohort RNA-seq analyses can yield unstable differential expression results, and underpowered designs are less likely to replicate well (Degen & Medo, 2025).

### 2.7 Gene Annotation and Functional Enrichment

Ensembl identifiers were mapped to HGNC symbols and biotypes using the Ensembl/GENCODE GRCh38 annotation. Accurate genome annotation is essential for RNA-seq quantification because annotations define gene coordinates, features, isoforms, and associated biotypes (Mudge *et al*., 2024). Stable gene identifiers are generally more reliable than symbols alone, because symbols and synonyms can be ambiguous (Juszczak *et al*., 2025). Over-representation analysis was performed separately for up- and down-regulated DEGs of each contrast with clusterProfiler against GO Biological Process, Cellular Component and Molecular Function, and KEGG pathways (Feu *et al*., 2023; Kanehisa *et al*., 2024). clusterProfiler provides enrichGO and enrichKEGG functions for GO- and KEGG-based enrichment analysis (Stead *et al*., 2025). GO is a comprehensive evidence-supported functional knowledge base with Biological Process, Molecular Function and Cellular Component branches (Feu *et al*., 2023; Aleksander *et al*., 2025). KEGG is a database resource for representing biological systems, and its pathway maps are a primary dataset for linking genes to molecular interaction networks (Zhang *et al*., 2023; Kanehisa *et al*., 2024). We considered terms with Benjamini–Hochberg-adjusted P < 0.05 enriched, consistent with common ORA practice (Stead *et al*., 2025; De Oliveira *et al*., 2026). The choice of background universe materially affects ORA results, and using the tested or annotated gene universe rather than an inappropriate default background is important for valid enrichment inference (Ziemann *et al*., 2024).

### 2.8 Visualisation

We produced PCA plots, volcano plots, and enrichment dot plots in R. Volcano-style displays of fold change against significance are standard in differential expression analysis (Stead *et al*., 2025; De Oliveira *et al*., 2026). clusterProfiler also supports visualisation of enrichment results, including dot-based summaries in which dot size and colour encode enrichment-related quantities (Fujita *et al*., 2024). Figures were numbered in order of citation; supplementary figures contained the remaining analyses.

## 3.0 Results

### 3.1 Cohort structure and global transcriptional variation

The cohort comprised 15 DLPFC donors: two Normal (A−T−), five pathology-ageing (A+T−) and eight AD (A+T+) cases (Table 1). Genotype was unevenly distributed across diagnoses: the two Normal donors were *APOE* E3/E3 and *TREM2* wild type, ε4 carriers were concentrated in PA and AD, and both E4/E4 donors were AD cases.

Principal component analysis of the donor-level expression profiles showed that the first two components explained 34% and 20% of the variance, respectively (Figure 1). The two Normal donors clustered together in the upper left of the plot; AD donors were spread along PC1, with the three most positive PC1 values (about 18–28) all belonging to AD cases, and four of the five PA donors lay at negative PC1 values with high dispersion along PC2. Diagnostic groups were therefore partially ordered along PC1 but not discretely separated, indicating that disease-associated variation coexists with substantial inter-individual heterogeneity.

**Figure 1.**
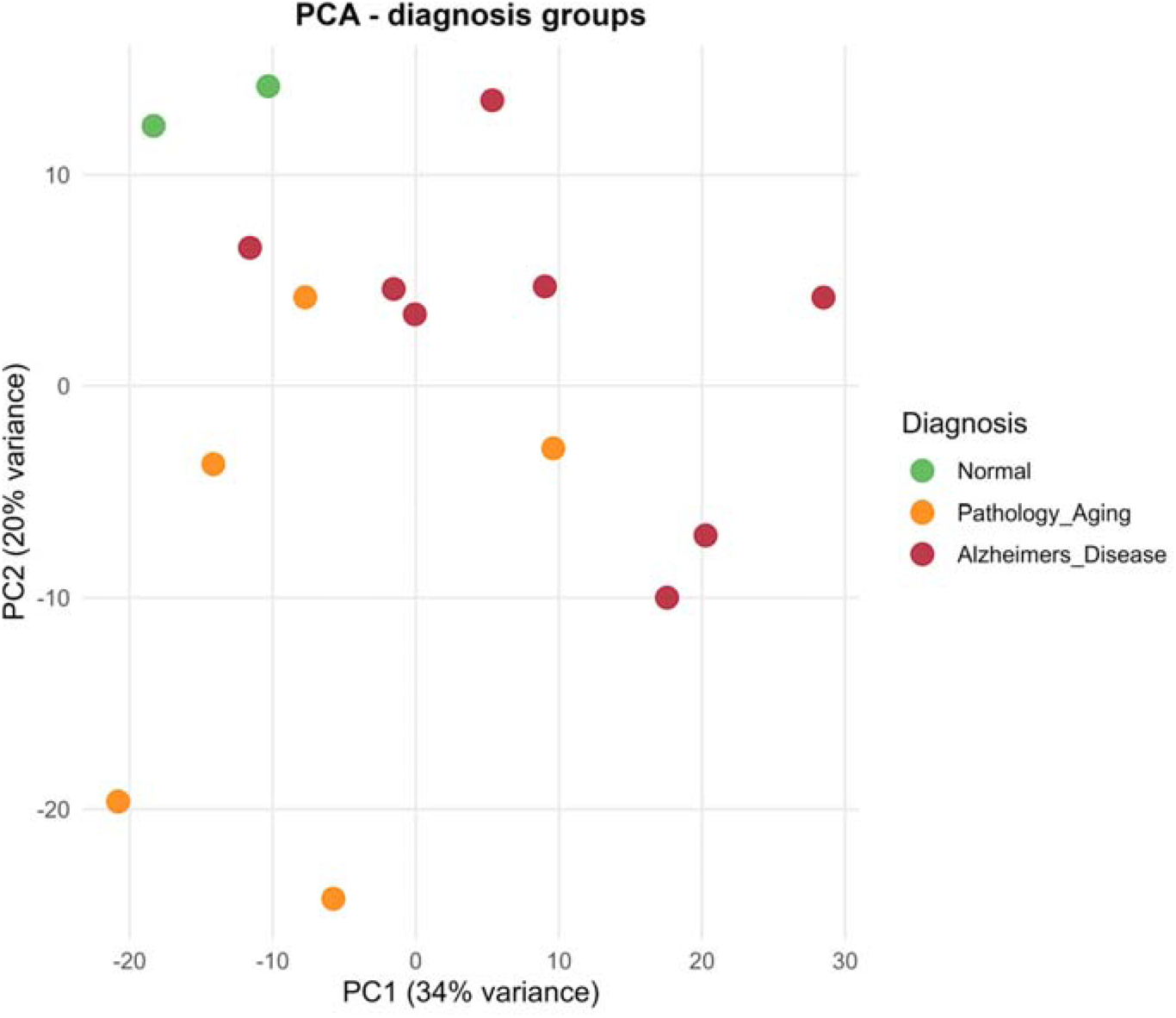
Principal component analysis of donor-level DLPFC transcriptomes. Each point is one donor (*n* = 15), coloured by neuropathological diagnosis (green, Normal; orange, pathology ageing; red, Alzheimer’s disease). PC1 and PC2 explain 34% and 20% of the variance, respectively. Groups are partially ordered along PC1, with considerable within-group dispersion.

### 3.2 Differential expression between Alzheimer’s disease and normal brain

Comparing AD with Normal donors, DESeq2 reported 406 genes with adjusted *P* < 0.05, of which 341 also met the effect-size criterion (|log FC| > 1): 223 were higher and 118 lower in AD (Table 2; Figure 2). Most DEGs were protein-coding (256; 75%), and 57 were long non-coding RNAs. The distribution of effect sizes was asymmetric, with a longer right tail of strongly induced genes (log FC up to 6.3 for *MS4A14*).

**Figure 2.**
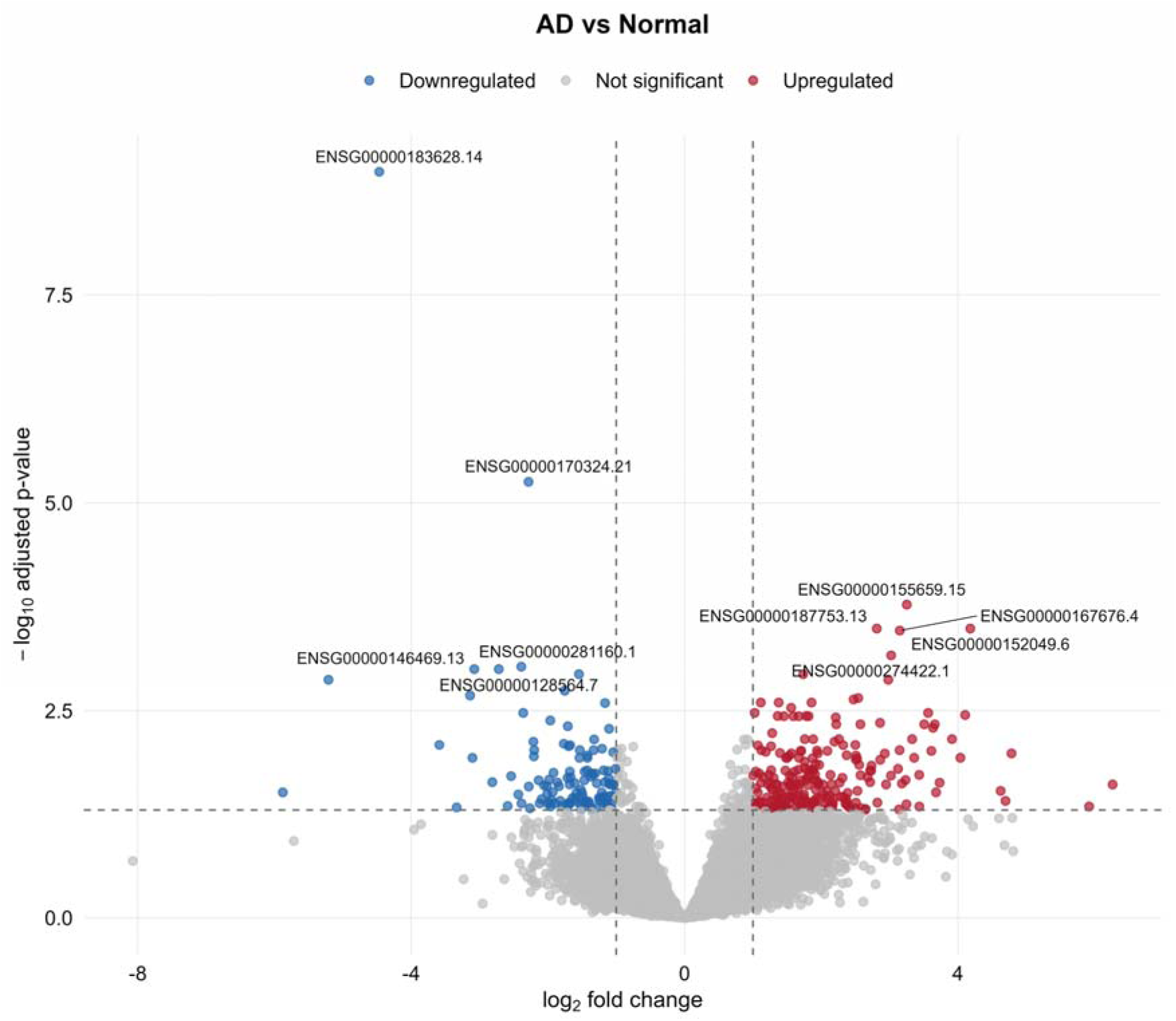
Volcano plot of differential expression, Alzheimer’s disease versus Normal. Each point is a gene; the *x*-axis shows log fold change (positive, higher in AD) and the *y*-axis shows −log adjusted *P*. Dashed lines mark adjusted *P* = 0.05 and |log FC| = 1. Red, up-regulated (*n* = 223); blue, down-regulated (*n* = 118); grey, not significant. Labelled points are the most significant genes (Ensembl IDs; corresponding symbols are given in Table 3).

**Table 2.** Summary of differential expression results for the three diagnostic contrasts.

| **Contrast** | **n vs n** | **Genes in results** | **Genes with adj. P assigned** | **adj. P < 0.05** | **DEGs (adj. P < 0.05, \|log□FC\| > 1)** | **Up** | **Down** | **Protein-coding** | **lncRNA** |
|---|---|---|---|---|---|---|---|---|---|
| AD vs Normal | 8 vs 2 | 37,558 | 21,517 | 406 | 341 | 223 | 118 | 256 | 57 |
| AD vs Pathology ageing | 8 vs 5 | 37,558 | 20,790 | 1089 | 496 | 358 | 138 | 313 | 136 |
| Pathology ageing vs Normal | 5 vs 2 | 37,558 | 28,057 | 6 | 6 | 5 | 1 | 5 | 1 |

For each pairwise diagnostic comparison, the number of genes carried into the DESeq2 model, the number for which an adjusted P value could be assigned after independent filtering, and the number of genes meeting the significance (adjusted P < 0.05) and effect-size (\|log□FC\| > 1) thresholds, with their direction and biotype. DEG, differentially expressed gene; FC, fold change. Positive log□FC denotes higher expression in the first-named group. Biotype counts refer to DEGs.

**Table 3.** Top differentially expressed genes, Alzheimer’s disease versus Normal.

| **Direction** | **Ensembl ID** | **Gene symbol** | **Biotype** | **Mean norm. count** | **log□FC** | **adj. P** |
|---|---|---|---|---|---|---|
| Up | ENSG00000155659 | *VSIG4* | protein coding | 245.5 | 3.25 | 1.7e-04 |
| Up | ENSG00000152049 | *KCNE4* | protein coding | 40.3 | 4.18 | 3.3e-04 |
| Up | ENSG00000187753 | *C9orf153* | protein coding | 65.0 | 2.81 | 3.3e-04 |
| Up | ENSG00000167676 | *PLIN4* | protein coding | 150.9 | 3.15 | 3.5e-04 |
| Up | ENSG00000274422 | — | lncRNA | 2862.1 | 3.02 | 6.9e-04 |
| Up | ENSG00000140443 | *IGF1R* | protein coding | 2085.8 | 1.74 | 0.001 |
| Up | ENSG00000145358 | *DDIT4L* | protein coding | 58.4 | 2.98 | 0.001 |
| Up | ENSG00000241170 | *RPL31P46* | transcribed processed pseudogene | 224.2 | 2.54 | 0.002 |
| Up | ENSG00000129810 | *SGO1* | protein coding | 46.8 | 2.47 | 0.002 |
| Up | ENSG00000103995 | *CEP152* | protein coding | 383.8 | 1.86 | 0.003 |
| Down | ENSG00000183628 | *DGCR6* | protein coding | 38.3 | −4.47 | 1.0e-09 |
| Down | ENSG00000170324 | *FRMPD2* | protein coding | 68.2 | −2.28 | 5.6e-06 |
| Down | ENSG00000281160 | — | lncRNA | 59.8 | −2.39 | 9.4e-04 |
| Down | ENSG00000128564 | *VGF* | protein coding | 726.0 | −2.72 | 0.001 |
| Down | ENSG00000146469 | *VIP* | protein coding | 252.7 | −3.08 | 0.001 |
| Down | ENSG00000204052 | *LRRC73* | protein coding | 210.6 | −1.55 | 0.001 |
| Down | ENSG00000147571 | *CRH* | protein coding | 199.2 | −5.21 | 0.001 |
| Down | ENSG00000265313 | *LINC01476* | lncRNA | 132.0 | −1.75 | 0.002 |
| Down | ENSG00000134443 | *GRP* | protein coding | 17.7 | −3.14 | 0.002 |
| Down | ENSG00000183828 | *NUDT14* | protein coding | 711.2 | −1.16 | 0.003 |

The ten most significant up-regulated and ten most significant down-regulated genes (ranked by adjusted P among genes with \|log□FC\| > 1) are listed with Ensembl identifier, gene symbol, biotype, mean normalised count and effect size. A dash indicates an Ensembl identifier that is absent from the annotation used for symbol assignment. Mean norm. count, DESeq2 baseMean (mean of size-factor-normalised counts across all donors). adj. P, Benjamini–Hochberg adjusted *P* value. log□FC > 0 indicates higher expression in AD.

The most statistically significant gene was *DGCR6* (log FC = −4.47; adjusted *P* = 1.0 × 10^−9^), followed by *FRMPD2* (log FC = −2.28; adjusted *P* = 5.6 × 10^−6^), both reduced in AD (Table 3; Figure 2). Among up-regulated genes, the highest-ranked included the complement-regulatory myeloid receptor *VSIG4* (log FC = 3.25; adjusted *P* = 1.7 × 10^−4^), *KCNE4*, *PLIN4* and *IGF1R*. Notably, the strongest down-regulated protein-coding genes with known neural functions were neuropeptides and secretory-granule proteins: *CRH* (log FC = −5.21), *GRP* (−3.14), *VIP* (−3.08) and *VGF* (−2.72).

### 3.3 Differential expression between Alzheimer’s disease and pathology ageing

To isolate changes associated with the transition from amyloid-only pathology to AD, we compared the eight AD donors with the five PA donors. This contrast produced the largest transcriptional response: 1,089 genes with adjusted *P* < 0.05 and 496 DEGs (358 up, 138 down; Table 2; Figure 3). The DEG set was enriched in non-coding transcripts relative to the AD-vs-Normal contrast (136 lncRNAs, 27%, versus 57, 17%).

**Figure 3.**
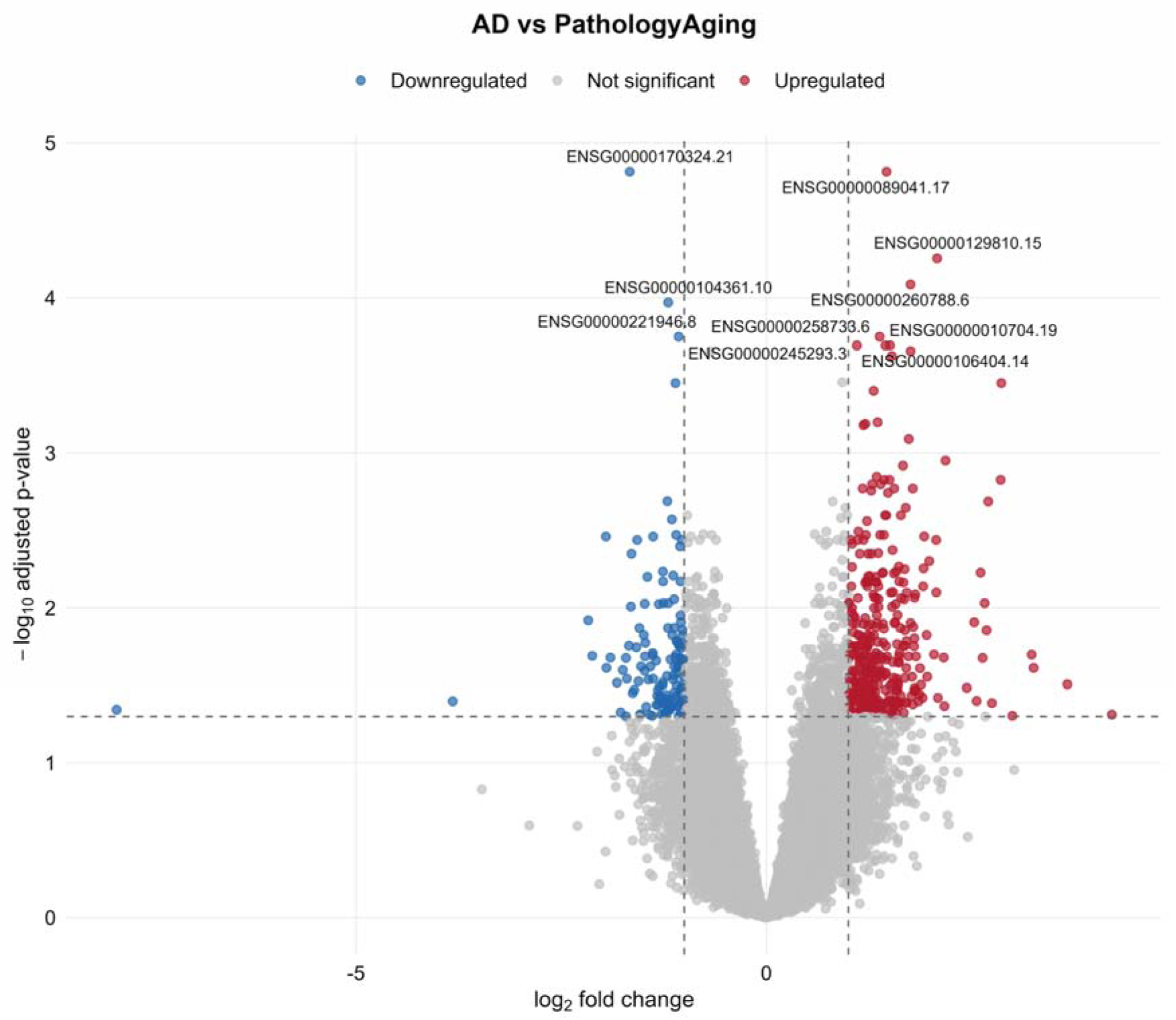
Volcano plot of differential expression, Alzheimer’s disease versus pathology ageing. Axes, thresholds and colours as in Figure 2. Positive log FC denotes higher expression in AD relative to PA. Red, up-regulated (*n* = 358); blue, down-regulated (*n* = 138). Labelled points are the most significant genes (Ensembl IDs; symbols in Table 4).

**Table 4.** Top differentially expressed genes, Alzheimer’s disease versus pathological ageing.

| Direction | Ensembl ID | Gene symbol | Biotype | Mean norm. count | log□FC | adj. P |
|---|---|---|---|---|---|---|
| Up | ENSG00000089041 | *P2RX7* | protein coding | 4475.7 | 1.46 | 1.5e-05 |
| Up | ENSG00000129810 | *SGO1* | protein coding | 46.8 | 2.08 | 5.6e-05 |
| Up | ENSG00000260788 | *CEDORA* | lncRNA | 2193.6 | 1.76 | 8.2e-05 |
| Up | ENSG00000258733 | *LINC02328* | lncRNA | 151.7 | 1.38 | 1.8e-04 |
| Up | ENSG00000010704 | *HFE* | protein coding | 71.2 | 1.51 | 2.0e-04 |
| Up | ENSG00000106404 | *CLDN15* | protein coding | 438.0 | 1.45 | 2.0e-04 |
| Up | ENSG00000245293 | *CYP2U1-AS1* | lncRNA | 617.9 | 1.10 | 2.0e-04 |
| Up | ENSG00000204161 | *TMEM273* | protein coding | 364.5 | 1.76 | 2.2e-04 |
| Up | ENSG00000261103 | — | lncRNA | 336.1 | 1.53 | 2.4e-04 |
| Up | ENSG00000260426 | — | lncRNA | 37.9 | 2.86 | 3.5e-04 |
| Down | ENSG00000170324 | *FRMPD2* | protein coding | 68.2 | −1.66 | 1.5e-05 |
| Down | ENSG00000104361 | *NIPAL2* | protein coding | 576.2 | −1.19 | 1.1e-04 |
| Down | ENSG00000221946 | *FXYD7* | protein coding | 2028.9 | −1.07 | 1.8e-04 |
| Down | ENSG00000189410 | *SH2D5* | protein coding | 367.3 | −1.11 | 3.5e-04 |
| Down | ENSG00000138100 | *TRIM54* | protein coding | 131.5 | −1.21 | 0.002 |
| Down | ENSG00000152034 | *MCHR2* | protein coding | 188.5 | −1.15 | 0.003 |
| Down | ENSG00000070159 | *PTPN3* | protein coding | 174.8 | −1.10 | 0.003 |
| Down | ENSG00000115828 | *QPCT* | protein coding | 213.2 | −1.38 | 0.003 |
| Down | ENSG00000146469 | *VIP* | protein coding | 252.7 | −1.95 | 0.003 |
| Down | ENSG00000146054 | *TRIM7* | protein coding | 93.0 | −1.03 | 0.004 |

The ten most significant up-regulated and ten most significant down-regulated genes in the AD-versus-PA contrast are shown, using the same ranking and column definitions as Table 3. This contrast isolates changes that distinguish symptomatic AD (A+T+) from amyloid-positive, tau-negative brains (A+T−). log□FC > 0 indicates higher expression in AD than in PA. A dash indicates an Ensembl identifier absent from the annotation used for symbol assignment.

The top up-regulated gene was *P2RX7* (log FC = 1.46; adjusted *P* = 1.5 × 10^−5^; mean normalised count 4,476), which encodes a purinergic receptor expressed predominantly by microglia, followed by *SGO1*, the lncRNA *CEDORA*, *HFE* and *CLDN15* (Table 4). Among down-regulated genes, *FRMPD2* (log FC = −1.66; adjusted *P* = 1.5 × 10^−5^), *NIPAL2*, *FXYD7* and *SH2D5* were the most significant, and the neuropeptide *VIP* (log FC = −1.95; adjusted *P* = 0.003) and neuropeptide receptor *MCHR2* were also reduced. The largest fold changes included *MZB1* and *IGKC* (immunoglobulin-related; log FC 4.2 and 3.7) up and *HLA-DRB5* (log FC = −7.9) down; these were of marginal significance (adjusted *P* 0.03–0.05) and should be treated cautiously.

### 3.4 Pathology aging versus normal brain

In stark contrast to the AD contrasts, only six genes were differentially expressed between PA and Normal donors (five up, one down; Table 5; Supplementary Figure S1). Four of these had clear roles in myeloid or metabolic biology: the macrophage/microglia receptors *VSIG4* (log FC = 3.57) and *CD163* (4.55), the insulin-like growth factor receptor *IGF1R* (1.95) and the glucocorticoid-metabolising enzyme *HSD11B2* (3.22). *DGCR6* was again the most significant gene and was reduced (log FC = −4.38). Strikingly, all six PA-vs-Normal DEGs were also DEGs in the AD-vs-Normal comparison, with the same direction and similar magnitude, indicating that these changes are already established in amyloid-positive, tau-negative brain and are not further amplified by tau pathology (for example, *VSIG4*, *CD163* and *IGF1R* did not differ between AD and PA).

**Table 5.** Differentially expressed genes, pathology aging versus Normal.

| Ensembl ID | Gene symbol | Biotype | Mean norm. count | log□FC | adj. P |
|---|---|---|---|---|---|
| ENSG00000183628 | *DGCR6* | protein coding | 38.3 | −4.38 | 1.2e-07 |
| ENSG00000155659 | *VSIG4* | protein coding | 245.5 | 3.57 | 8.4e-05 |
| ENSG00000177575 | *CD163* | protein coding | 293.8 | 4.55 | 2.1e-04 |
| ENSG00000140443 | *IGF1R* | protein coding | 2085.8 | 1.95 | 8.5e-04 |
| ENSG00000176387 | *HSD11B2* | protein coding | 113.5 | 3.22 | 0.009 |
| ENSG00000271967 | — | lncRNA | 56.8 | 1.85 | 0.034 |

All six genes that met the significance and effect-size criteria in the pathology-aging-versus-Normal contrast are listed with their annotation, expression level and effect size. They represent the earliest detectable transcriptional changes in the A/T continuum in this dataset. log□FC > 0 indicates higher expression in PA. The unannotated lncRNA (ENSG00000271967) is absent from the annotation used for symbol assignment.

### 3.5 Overlap between contrasts and concordance of effects

Fifty genes were DEGs in both AD contrasts (AD vs Normal and AD vs PA), and all 50 changed in the same direction (22 up, 28 down), a set that is unlikely to reflect group-specific noise in either comparison. The shared down-regulated genes featured neuropeptides and their receptors (*VIP*, *VGF*, *GRP*, *TAC1*, *TACR2*, *MCHR1*, *MCHR2*, *PCSK1*, *RBP4*) together with *FRMPD2*, *PPP1R11*, *NSG1* and *SLC32A1*, the vesicular GABA transporter. Shared up-regulated genes included *TLR4*, *PARP9*, *PECAM1*, *KCNE4* and *C9orf153*. By contrast, the complement and macrophage genes that were elevated in AD versus Normal (*C1QA*, *C1QB*, *C1QC*, *VSIG4*, *CD163*) were not different between AD and PA (|log FC| ≤ 1), mirroring their early rise in PA. Neuropeptide transcripts showed a graded pattern: for example, *VIP* log FC was −1.12 in PA vs Normal (not significant), −1.95 in AD vs PA and −3.08 in AD vs Normal, and *VGF* log FC was −1.52, −1.20 and −2.72, respectively.

### 3.6 Functional enrichment of genes altered in Alzheimer’s disease versus normal brain

Genes up-regulated in AD relative to Normal were enriched in 35 GO BP terms, dominated by immune processes (Table 6; Figure 4A). The top terms included response to type II interferon (8 genes, adjusted *P* = 0.016), myeloid leukocyte activation (10 genes), leukocyte-mediated immunity (13 genes), adaptive immune response (14 genes) and complement activation (5 genes; *VSIG4*, *C1QA*, *C1QB*, *C1QC*, *CFH*; fold enrichment 11.6). Synapse pruning (*C1QA*, *C1QB*, *C1QC*) was also enriched (fold enrichment 20.9). At the cellular component level, the collagen-containing extracellular matrix (13 genes) was enriched; at the molecular-function level, Wnt receptor activity (*FZD1*, *FZD8*, *FZD9*, *LRP5*; fold enrichment 32.2, adjusted *P* = 0.002) and immune receptor activity (Figure 4B; Supplementary Figure S5). The single enriched KEGG pathway was *Staphylococcus aureus* infection (adjusted *P* = 0.003), which in this context reflects the complement and Fcγ-receptor genes (*C1QA*, *C1QB*, *C1QC*, *FCGR2A*, *FCGR3A*, *CFH*) rather than a microbial signature.

**Figure 4.**
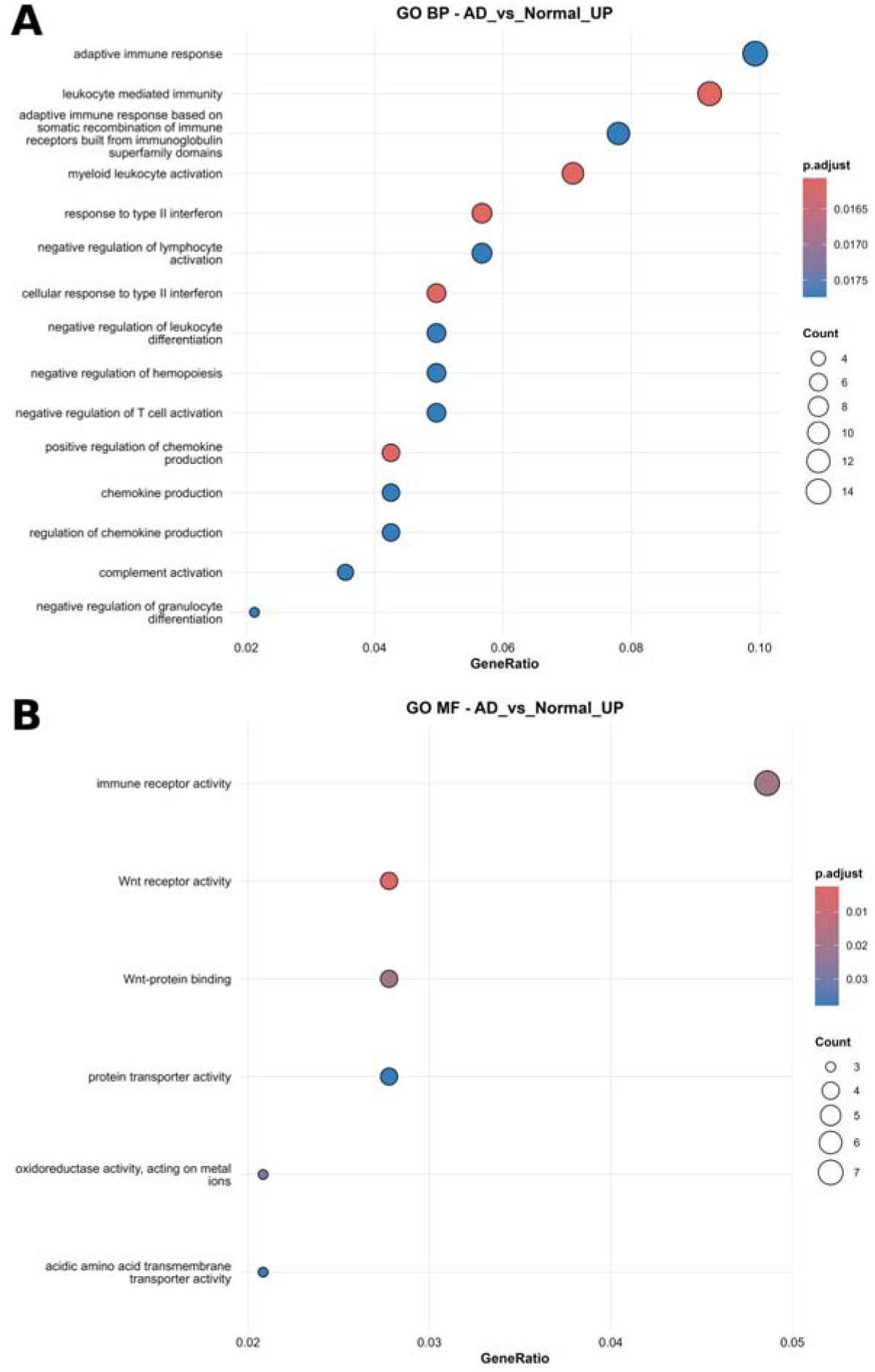
GO enrichment of genes up-regulated in Alzheimer’s disease versus Normal. (A) Biological process; (B) molecular function. Gene ratio is shown on the *x*-axis, point size indicates the number of genes in the term, and colour indicates adjusted *P*. Immune activation (interferon-γ response, myeloid activation, adaptive immunity, complement) dominates the biological-process results, and Wnt/Fzd receptor activity and immune receptor activity dominate molecular function. GO CC and KEGG results are in Supplementary Figure S5.

**Table 6.**
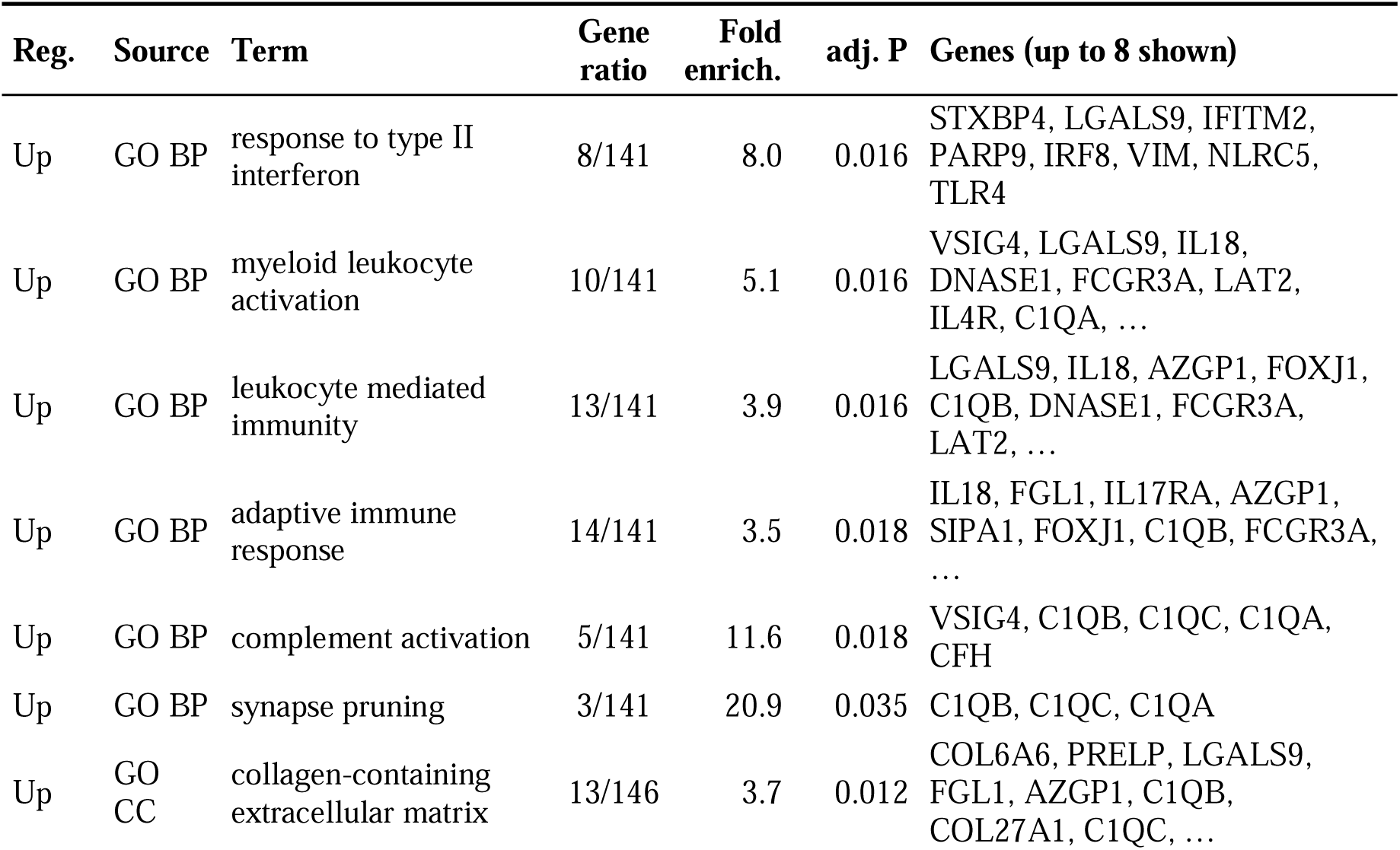
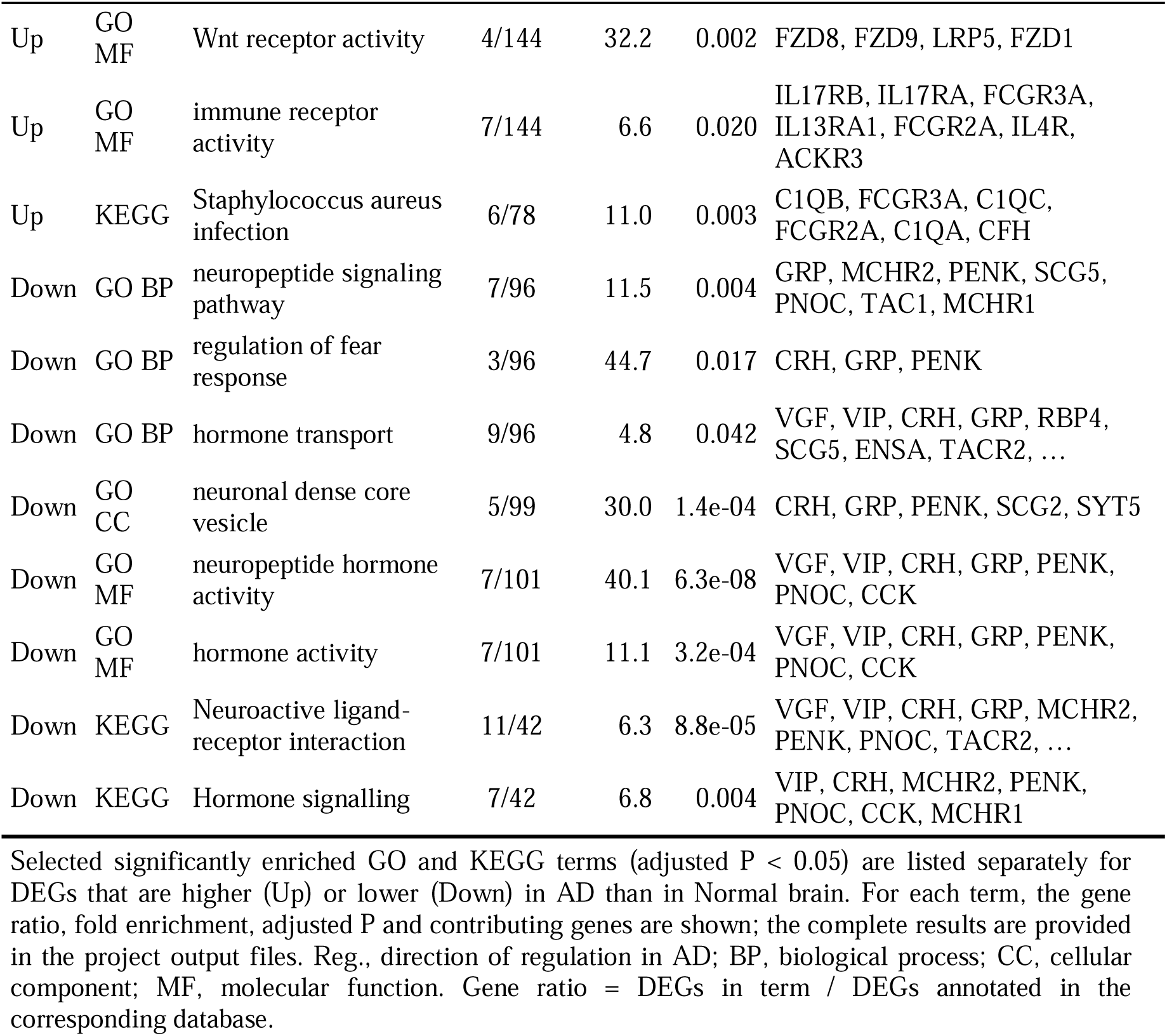
Functional enrichment of up- and down-regulated genes, Alzheimer’s disease versus Normal.

Genes **down-regulated** in AD versus Normal were enriched in a compact set of neuronal-secretory functions (Table 6; Figure 5). Neuropeptide signalling pathway (7 genes: *GRP*, *MCHR1*, *MCHR2*, *PENK*, *PNOC*, *SCG5*, *TAC1*; fold enrichment 11.5, adjusted *P* = 0.004), regulation of the fear response (*n* = 3; fold enrichment 44.7) and hormone transport (9 genes: *VGF*, *VIP*, *CRH*, *GRP*, *RBP4*, *SCG5*, *ENSA*, *TACR2*, *CRYM*) were enriched, together with the neuronal dense core vesicle (5 genes; adjusted *P* = 1.4 × 10^−4^) and neuropeptide hormone activity (7 genes; fold enrichment 40.1; adjusted *P* = 6.3 × 10^−8^). KEGG analysis identified neuroactive ligand–receptor interaction (11 of 42 genes; adjusted *P* = 8.8 × 10^−5^) and hormone signalling (adjusted *P* = 0.004; Figure 5B). The remaining cellular-component and molecular-function terms are in Supplementary Figure S6.

**Figure 5.**
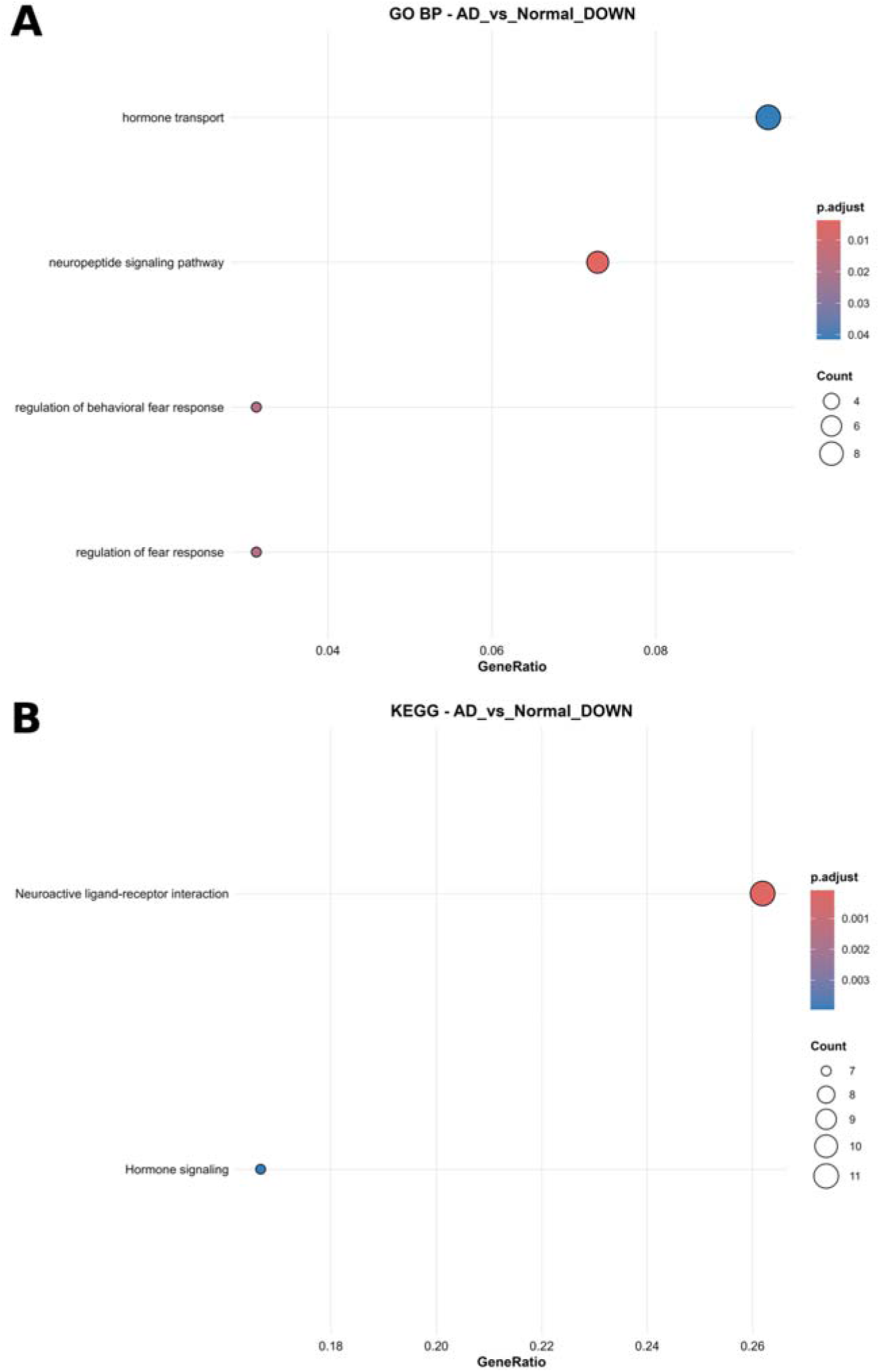
GO and KEGG enrichment of genes down-regulated in Alzheimer’s disease versus Normal. (A) GO biological process; (B) KEGG pathways. Neuropeptide signalling and neuroactive ligand–receptor interaction are the principal enriched themes. Plot encoding as in Figure 4. GO CC and MF results are in Supplementary Figure S6.

### 3.7 Functional enrichment of genes altered in Alzheimer’s disease versus pathology aging

Genes **up-regulated** in AD relative to PA were strongly enriched in glial and myelination biology (Table 7; Figure 6A). Gliogenesis (21 of 195 genes; fold enrichment 5.1; adjusted *P* = 2.6 × 10^−6^), glial cell differentiation, myelination (12 genes; adjusted *P* = 3.6 × 10^−4^), oligodendrocyte differentiation (10 genes: *BOK*, *QKI*, *CNTN2*, *MYRF*, *SOX10*, *FA2H*, *CXCR4*, *DAAM2*, *DUSP10*, *SLC45A3*; adjusted *P* = 4.3 × 10^−4^) and myelin maintenance (*MYRF*, *SH3TC2*, *FA2H*, *CXCR4*) were enriched among 18 significant BP terms. The only other significant terms were the cortical actin cytoskeleton (CC) and two MF terms, myosin II binding (*LLGL1*, *LLGL2*, *GSN*, *SHROOM4*) and mechanosensitive ion channel activity (*PIEZO2*, *TMC6*, *TMC7*, *TMEM63A*) (Figure 6B; Supplementary Figure S7A). No KEGG pathway was enriched among the up-regulated genes.

**Figure 6.**
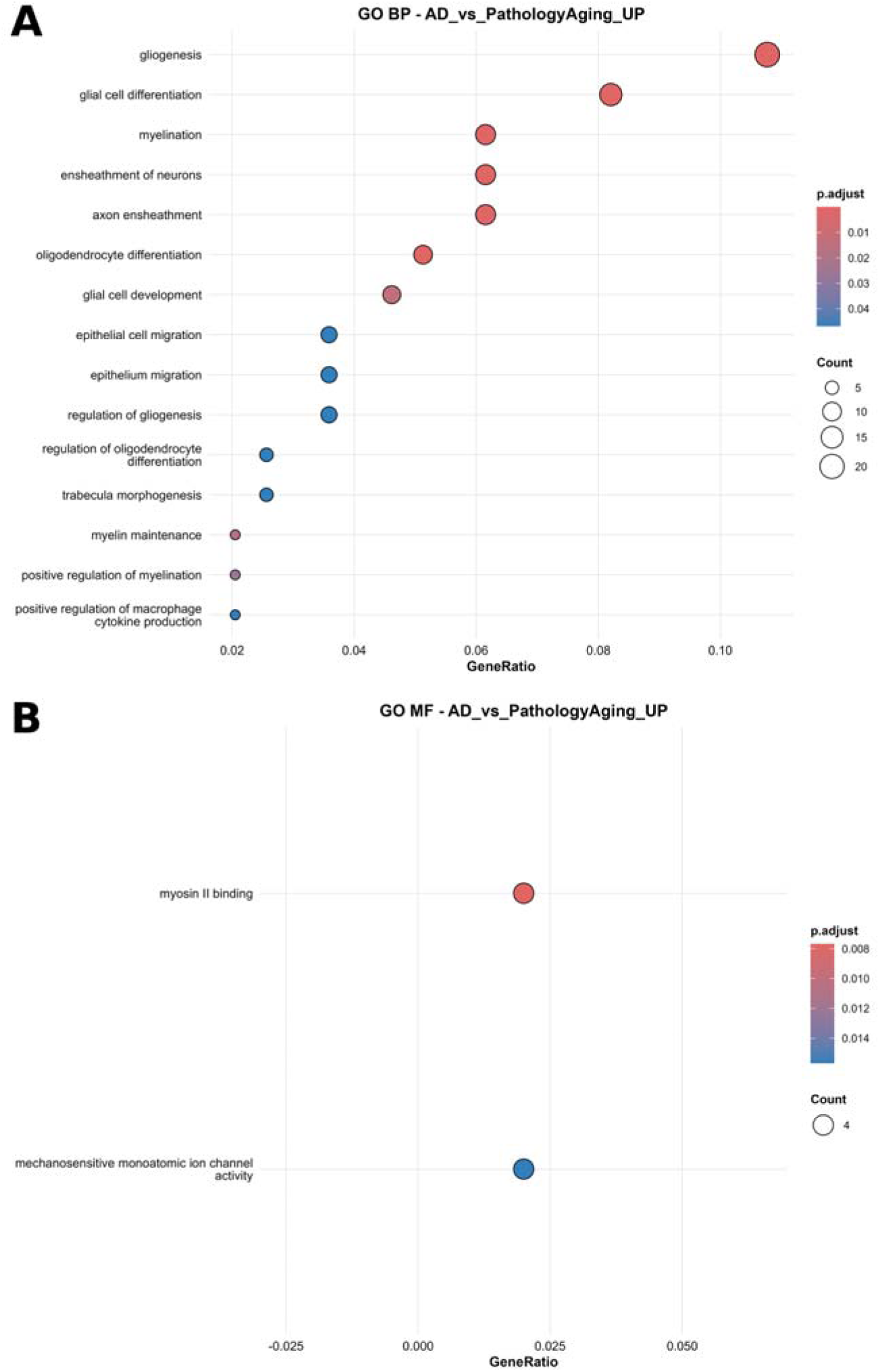
GO enrichment of genes up-regulated in Alzheimer’s disease versus pathology aging. (A) Biological process, showing gliogenesis, myelination and oligodendrocyte differentiation; (B) molecular function. Plot encoding as in Figure 4. The GO CC result (cortical actin cytoskeleton) is shown in Supplementary Figure S7A.

**Table 7.** Functional enrichment of up- and down-regulated genes, Alzheimer’s disease versus pathology aging.

| Reg. | Source | Term | Gene ratio | Fold enrich. | adj. P | Genes (up to 8 shown) |
|---|---|---|---|---|---|---|
| Up | GO BP | Gliogenesis | 21/195 | 5.1 | 2.6e-06 | BOK, QKI, CSF1, ARHGEF10, CNTN2, TLR4, GJC2, LRP2, … |
| Up | GO BP | Myelination | 12/195 | 6.3 | 3.6e-04 | CDK18, UGT8, QKI, ARHGEF10, CNTN2, BCAS1, MYRF, SH3TC2, … |
| Up | GO BP | oligodendrocyte differentiation | 10/195 | 7.5 | 4.3e-04 | BOK, QKI, CNTN2, MYRF, SOX10, FA2H, CXCR4, DAAM2, … |
| Up | GO BP | myelin maintenance | 4/195 | 19.0 | 0.018 | MYRF, SH3TC2, FA2H, CXCR4 |
| Up | GO BP | central nervous system myelination | 4/195 | 12.4 | 0.047 | CNTN2, MYRF, SOX10, FA2H |
| Up | GO CC | cortical actin cytoskeleton | 7/205 | 7.2 | 0.016 | PIEZO2, MYO1E, LLGL2, ANLN, SHROOM4, GSN, LLGL1 |
| Up | GO MF | myosin II binding | 4/200 | 24.9 | 0.008 | LLGL2, SHROOM4, GSN, LLGL1 |
| Up | GO MF | mechanosensitive monoatomic ion channel activity | 4/200 | 18.0 | 0.016 | PIEZO2, TMC6, TMC7, TMEM63A |
| Down | GO BP | regulation of sodium ion transmembrane transport | 5/86 | 19.9 | 0.008 | FXYD7, PTPN3, TESC, FGF12, GRP |
| Down | GO MF | growth factor activity | 6/91 | 7.8 | 0.016 | FGF9, NRG4, FGF12, THPO, BMP4, VGF |
| Down | GO MF | neuropeptide hormone activity | 3/91 | 19.1 | 0.026 | VIP, VGF, GRP |
| Down | KEGG | Neuroactive ligand-receptor interaction | 9/34 | 6.4 | 6.1e-04 | MCHR2, VIP, MCHR1, TACR2, TAC1, KISS1R, HTR6, VGF, … |

Selected significantly enriched GO and KEGG terms are shown for DEGs higher (Up) or lower (Down) in AD than in pathology-aging brain, with the same column definitions as Table 6. These terms characterise the programmes that distinguish tau-positive AD from amyloid-positive, tau-negative brain.

Genes **down-regulated** in AD versus PA were again enriched for neuropeptide-related biology (Table 7; Figure 7), but also for ion-transport regulation. Regulation of sodium ion transmembrane transport (*FXYD7*, *PTPN3*, *TESC*, *FGF12*, *GRP*; fold enrichment 19.9; adjusted *P* = 0.008), growth-factor activity (*FGF9*, *NRG4*, *FGF12*, *THPO*, *BMP4*, *VGF*), neuropeptide hormone activity (*VIP*, *VGF*, *GRP*) and, again, KEGG neuroactive ligand–receptor interaction (9 of 34 genes; adjusted *P* = 6.1 × 10^−4^) were enriched. No cellular-component term reached significance (Supplementary Figure S7B shows the MF dot plot).

**Figure 7.**
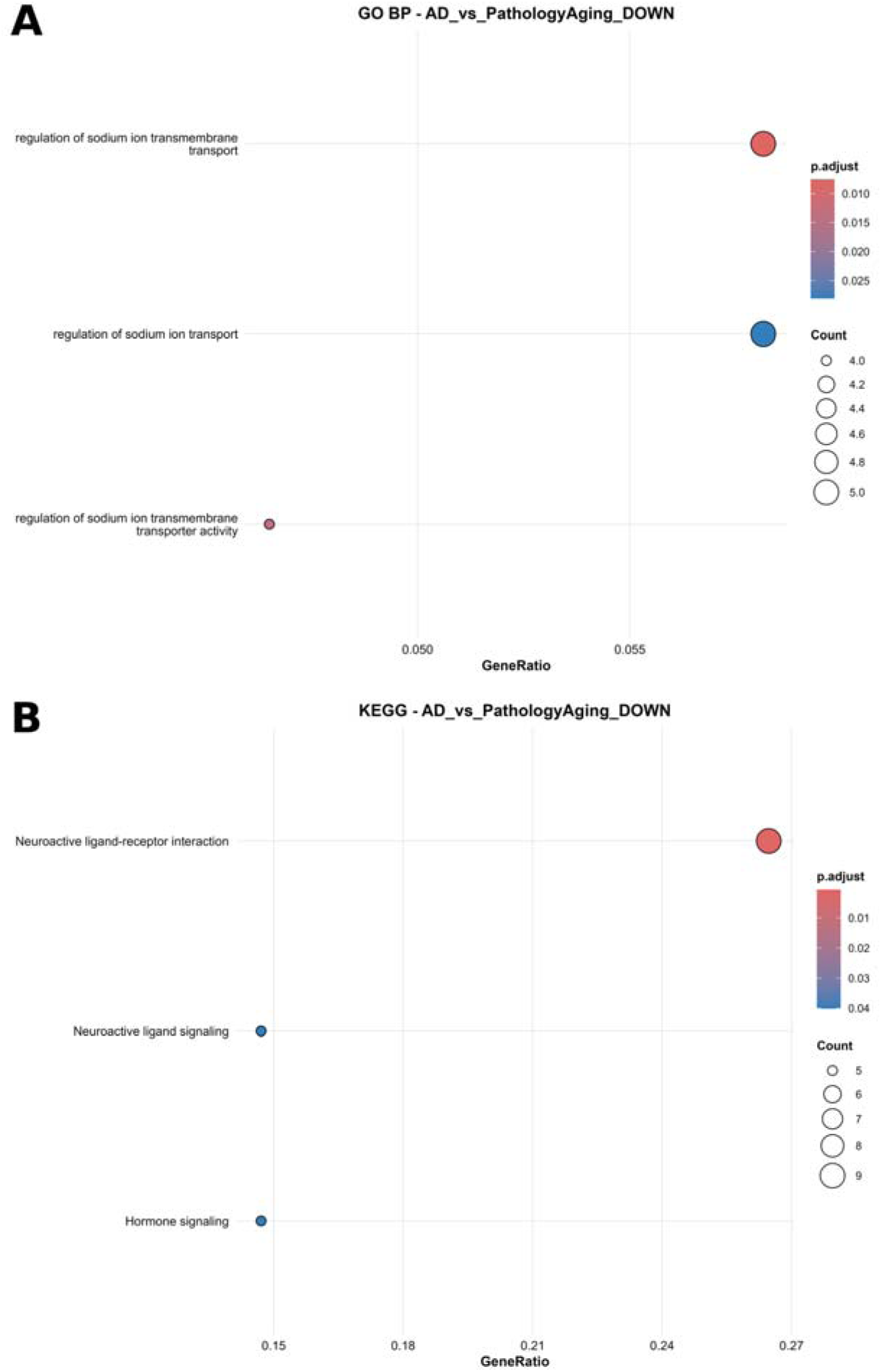
GO and KEGG enrichment of genes down-regulated in Alzheimer’s disease versus pathology aging. (A) GO biological process (regulation of sodium ion transmembrane transport); (B) KEGG pathways (neuroactive ligand–receptor interaction, neuroactive ligand signalling, hormone signalling). Plot encoding as in Figure 4.

### 3.8 Enrichment among pathology-aging versus normal DEGs

Because only six genes were differentially expressed between PA and Normal, enrichment results for this contrast rest on four or five annotated genes and are statistically fragile (Table 8; Supplementary Figure S8). Among up-regulated genes, 87 BP and 47 MF terms reached adjusted *P* < 0.05, but nearly all were driven by one or two genes (gene ratios of 1/4 or 2/4), notably *IGF1R* and *HSD11B2*; they include response to glucocorticoid, response to insulin, complement binding (*VSIG4*) and scavenger receptor activity (*CD163*). No down-regulated term and no KEGG pathway was enriched. We therefore regard this analysis as descriptive and do not draw inferences from individual terms.

**Table 8.** Enrichment of up-regulated genes, pathology aging versus Normal.

| **Reg.** | **Source** | **Term** | **Gene ratio** | **Fold enrich.** | **adj. P** | **Genes (up to 8 shown)** |
|---|---|---|---|---|---|---|
| Up | GO BP | response to glucocorticoid | 2/4 | 60.6 | 0.046 | IGF1R, HSD11B2 |
| Up | GO BP | complement activation, alternative pathway | 1/4 | 328.1 | 0.046 | VSIG4 |
| Up | GO BP | response to insulin | 2/4 | 25.5 | 0.046 | IGF1R, HSD11B2 |
| Up | GO MF | insulin-like growth factor binding | 1/4 | 213.3 | 0.020 | IGF1R |
| Up | GO MF | complement binding | 1/4 | 225.2 | 0.020 | VSIG4 |
| Up | GO MF | scavenger receptor activity | 1/4 | 168.9 | 0.020 | CD163 |

Representative GO terms enriched among the five genes up-regulated in pathology aging relative to Normal. The gene ratios (numerator of one or two genes) show that each term rests on very few genes, so the table should be read as descriptive rather than inferential. No term was enriched among down-regulated genes, and no KEGG pathway was enriched in either direction. Abbreviations as in Table 6.

### 3.9 Exploratory genotype-stratified analyses

The *APOE* E4/E4 versus E3/E3 contrast produced 235 DEGs (50 up, 185 down; Table 9; Supplementary Figure S2), including reduced expression of the glycolytic genes *HK1* and *ALDOC*, the NMDA receptor subunit *GRIN2B* and *TCEAL5*. However, both E4/E4 donors were AD cases while three of the five E3/E3 donors were PA or Normal (Table 1), so this contrast cannot be separated from diagnosis. The E3/E4 versus E3/E3 contrast produced seven DEGs (Supplementary Figure S3) and the *TREM2* R47H versus wild-type contrast two (Supplementary Figure S4); all of these were pseudogenes or lncRNAs, and most had mean normalised counts near one with implausibly large log FC (|log FC| ≈ 17–21), the typical signature of low-count artefacts. Neither *APOE* nor *TREM2* transcripts themselves differed significantly between diagnostic groups (AD vs Normal: *APOE* log FC = 0.81, adjusted *P* = 0.38; *TREM2* log FC = 1.45, adjusted *P* = 0.33).

**Table 9.** Exploratory genotype-stratified differential expression.

| Contrast | n vs n | DEGs | Up | Down | Interpretation |
|---|---|---|---|---|---|
| *APOE* E4/E4 vs E3/E3 | 2 vs 5 | 235 | 50 | 185 | Both E4/E4 donors are AD cases and three of five E3/E3 donors are non-AD; confounded with diagnosis. Reduced: *HK1*, *ALDOC*, *GRIN2B*, *TCEAL5*. |
| *APOE* E3/E4 vs E3/E3 | 8 vs 5 | 7 | 1 | 6 | Non-coding / pseudogene signals; five of seven have mean normalised counts < 1.5 (likely low-count artefacts). |
| *TREM2* p.Arg47His vs wild type | 4 vs 11 | 2 | 2 | 0 | Two processed pseudogenes with mean normalised counts ≈ 1 (likely artefacts); no coding-gene signal. |

This table summarises the secondary contrasts for *APOE* and *TREM2* genotype, giving the group sizes, DEG counts and an interpretation of each result in light of confounding with diagnosis and the low expression of the leading genes. DEGs, adjusted *P* < 0.05 and |log□FC| > 1. These contrasts were not adjusted for diagnosis and are hypothesis-generating only.

## 4.0 Discussion

Using DLPFC RNA-seq from donors stratified by amyloid and tau pathology, we found that the transcriptional response in AD is dominated by three themes: (i) a depletion of neuropeptide and hormone-signalling transcripts that is already directionally evident in pathology aging and is amplified in AD; (ii) an activation of interferon-γ, myeloid and complement programmes that is established by the pathology-aging stage and is not further amplified by tau pathology; and (iii) an AD-specific induction of gliogenesis, oligodendrocyte differentiation and myelination genes and of the microglial receptor P2RX7. These results indicate that the A−T−→A+T−→A+T+ continuum is not a single monotonic transcriptional gradient, but comprises distinct early immune and later neuronal/glial components (Gazestani *et al*., 2023; Gabitto *et al*., 2024).

The most consistent neuronal signature was reduced expression of neuropeptides and their receptors in AD: VGF, VIP, CRH, GRP, TAC1, PENK, SCG5, CCK, PNOC, and MCHR1/MCHR2 and TACR2. Many such transcripts are characteristic of vulnerable neuronal populations, including inhibitory interneuron subclasses that are depleted in AD, making enrichment of dense-core vesicle and neuropeptide-related terms biologically plausible (Mathys *et al*., 2023; Gabitto *et al*., 2024; Mathys *et al*., 2024). Of particular interest, VGF has emerged as a reproducible AD-linked neuronal factor: it was highlighted as a top regulator of AD-associated networks in a large postmortem multi-omics study and is reduced across the AD continuum in CSF, including preclinical stages (Lu *et al*., 2025). Independent synthesis of bulk RNA-seq studies also found VGF downregulated in both frontal and temporal cortex, supporting its relevance beyond a single cohort (Fox *et al*., 2025). The shared AD-vs-Normal and AD-vs-PA neuropeptide losses in our data therefore fit a broader literature in which synaptic and neurotrophic signaling genes decline as pathology advances, rather than reflecting amyloid positivity alone (Mathys *et al*., 2023; Nakatsuka *et al*., 2025). This interpretation is also compatible with evidence that early cortical amyloid responses can include transient neuronal states before later-stage neuronal depletion, so bulk RNA-seq may capture a mixture of changing cell states and cell loss (Gazestani *et al*., 2023; Mathys *et al*., 2024).

Immune genes were the main up-regulated signal in AD versus Normal: interferon-γ response, myeloid leukocyte activation, adaptive immunity, chemokine production and complement activation, with C1QA, C1QB and C1QC at the centre of several terms. This aligns with multiple single-cell and multimodal studies showing that early AD-associated microglial changes include inflammatory programs, interferon-response genes, antigen-presentation genes, and complement components such as C1QA and C1QB (Gazestani *et al*., 2023; Gabitto *et al*., 2024; Nakatsuka *et al*., 2025). C1q-mediated synapse elimination is mechanistically plausible because complement overactivation drives microglial synapse loss in neurodegeneration, and experimental blockade or deletion of C1q reduces microglial synaptic engulfment and improves functional outcomes (Chung *et al*., 2023; Wilton *et al*., 2023; Zhou *et al*., 2023; Petrisko *et al*., 2026). Complement proteins are also elevated and localized to plaques and tangles in AD brain, strengthening the relevance of our synapse-pruning enrichment result (Batista *et al*., 2024).

Several features of our data add nuance. Early-stage studies support the view that amyloid-positive cortex already shows microglial activation and inflammatory transcriptional remodeling, often before heavy tau burden or overt neuronal loss (Gazestani *et al*., 2023; Gabitto *et al*., 2024). Early upregulation of plaque-induced and complement genes, including C1qa/C1qb, has also been observed in pseudoprogression analyses and spatial plaque-proximal microglia (Gabitto *et al*., 2024; Mary *et al*., 2024). That pattern is consistent with our observation that VSIG4, CD163, IGF1R, and directionally increased C1q genes are already evident in pathology aging, although the broader literature does not establish that these programs necessarily plateau after tau emerges; rather, several studies report similar but stronger microglial perturbations with increasing pathological burden (Gazestani *et al*., 2023; Mancuso *et al*., 2024). A more cautious interpretation is therefore that myeloid-complement activation appears early in amyloid-positive brain and remains prominent with progression (Gabitto *et al*., 2024; Gazestani *et al*., 2023).

The link between amyloid, glial activation, and downstream neurodegeneration is supported by work positioning microglia as a bridge between amyloid deposition and later tau-linked injury (Bivona *et al*., 2023). AD genetics also converges on microglial and immune pathways, with AD-risk genes repeatedly enriched in disease-associated microglial signatures (Brase *et al*., 2023; Gazestani *et al*., 2023; Nakatsuka *et al*., 2025). TLR4 upregulation in our AD contrasts is compatible with the broader literature implicating microglial TLR4/NLRP3 signaling in AD neuroinflammation (Li *et al*., 2023; Long *et al*., 2024). Likewise, the AD-specific increase in P2RX7 is biologically plausible because P2X7 receptors are among the receptors linked to NLRP3 inflammasome activation in AD (Maurya *et al*., 2025; McGroarty *et al*., 2025). Because bulk tissue conflates regulation within cells with shifts in cell abundance, increased myeloid transcripts may reflect both activated resident microglia and changes in microglial/macrophage composition, consistent with reports of multiple AD-related microglial states and declining homeostatic markers during progression (Gabitto *et al*., 2024; Mancuso *et al*., 2024).

Relative to PA, AD brains displayed up-regulation of genes governing gliogenesis, oligodendrocyte differentiation and myelination, including SOX10, MYRF, FA2H, and UGT8. This fits a growing literature implicating oligodendrocyte-lineage remodeling in AD and identifying disease-related oligodendrocyte states, altered myelination programs, and region-specific remyelination responses (Brase *et al*., 2023; Ziar *et al*., 2025). Multimodal pseudoprogression analyses further suggest that an early AD phase includes oligodendrocyte loss together with increased OPC differentiation and remyelination factors, consistent with a compensatory response (Gabitto *et al*., 2024). MYRF is directly relevant here because it regulates myelination and is increased early in oligodendrocyte responses along AD pseudoprogression trajectories (Gabitto *et al*., 2024). Human and animal studies also support the broader premise that myelin dysfunction is an early event in AD and can actively worsen pathology rather than simply reflect late degeneration (Depp *et al*., 2023; Zota *et al*., 2024; Ziar *et al*., 2025).

At the same time, the timing of oligodendrocyte-lineage changes is complex. Some studies place remyelination-related responses in early amyloid-associated phases rather than only after tau-positive transition (Gabitto *et al*., 2024), whereas others emphasize later downregulation of myelin genes and broader oligodendrocyte dysfunction as pathology becomes severe (Gabitto *et al*., 2024). Our finding of stronger oligodendrocyte-lineage induction in AD versus PA therefore appears compatible with stage-dependent remodeling, but it should not be taken to mean that such programs are absent from earlier disease biology (Gazestani *et al*., 2023; Ziar *et al*., 2025). Experimental work also supports a causal connection between myelin dysfunction, amyloid deposition, and altered microglial states, reinforcing the relevance of this axis (Depp *et al*., 2023; Zota *et al*., 2024). The enrichment of mechanosensitive-channel genes among AD-up genes is more speculative, although age-related mechanical changes in the OPC niche have been proposed to impair OPC function, making this a plausible but still hypothesis-generating signal (Zou *et al*., 2023).

Only six genes distinguished PA from Normal brain, y*et al*l six were also altered in AD with the same direction, so they may mark the earliest detectable transcriptional footprint of amyloid pathology in this cohort. The small number of PA-vs-Normal DEGs should not be interpreted as absence of biology, because underpowered RNA-seq studies often show poor recall and low replicability even when some true signals are present (Degen & Medo, 2025). Small-N bulk RNA-seq designs are especially vulnerable to unstable effect-size estimates and misleading false positives, and raising fold-change thresholds does not solve that problem (Halasz *et al*., 2025). That caveat is directly relevant here, given the two-control design. The enrichment results for PA-vs-Normal should also be treated cautiously because pathway methods can generate unstable or false-positive results in contrasts with few or no DEGs (Candia & Ferrucci, 2024; Stead *et al*., 2025).

We detected no robust APOE-or TREM2-specific signal. That negative result should be interpreted cautiously because APOE and TREM2 effects are often cell-state-specific rather than readily visible in bulk abundance measures (Brase *et al*., 2023). In single-nucleus data, APOE ε4 and TREM2 risk variation is associated with distinct transcriptional states, including ferroptosis-like signatures in inhibitory neurons and autophagy-lysosomal dysregulation in TREM2-linked oligodendrocytes (Brase *et al*., 2023). Human microglia xenograft work likewise shows that APOE polymorphisms and TREM2 R47H differentially modulate microglial state transitions in response to amyloid pathology (Mancuso *et al*., 2024). The absence of APOE or TREM2 transcript-level differences in our bulk data is therefore consistent with functional effects driven by variant biology and cell-state context rather than by total mRNA abundance (Brase *et al*., 2023; Mancuso *et al*., 2024).

The reduced glycolytic genes in the E4/E4 contrast are biologically interesting because APOE4 is linked to metabolic dysfunction and impaired glial handling of lipids and glucose-related pathways ((Jung *et al*., 2025). Cerebral glucose metabolism declines in AD and is strongly linked to cognitive impairment (Jung *et al*., 2025). Oligodendrocyte glycolytic and ketolytic pathways have also been reported as selectively damaged in AD brain tissue. But because both E4/E4 donors in this cohort are AD cases, the present contrast cannot disentangle genotype from diagnosis.

Strengths of this study include the A/T-based group definition, use of a single cortical region, inclusion of pathology aging as an intermediate state, and explicit concordance analysis across contrasts. Several limitations remain important. With 15 donors and only two controls, statistical power is low and reproducibility is limited in principle (Degen & Medo, 2025; Halasz *et al*., 2025). Bulk tissue averages across neurons, glia, and vascular cells, so transcript shifts can arise from compositional changes as well as within-cell regulation (Gabitto *et al*., 2024; Mathys *et al*., 2024). Single-cell and single-nucleus atlases now show that AD involves multiple microglial, neuronal, and oligodendrocyte states that are obscured in homogenate data (Gazestani *et al*., 2023; Mancuso *et al*., 2024; Ziar *et al*., 2025). Independent validation in cohorts such as ROSMAP and Mount Sinai Brain Bank is feasible and desirable because these resources have supported cross-cohort transcriptomic analyses and replication efforts in AD (Fox *et al*., 2025; Gupta *et al*., 2025; Aksu *et al*., 2026).

The graded neuropeptide signature and the AD-specific glial/myelination programme are plausible candidates for tracking disease stage and for therapeutic follow-up. The strongest convergent signal is that amyloid-positive brain already shows inflammatory microglial and complement activation, while later progression is accompanied by additional neuronal loss signatures and oligodendrocyte-lineage remodeling (Gabitto *et al*., 2024).

## 5.0 Conclusions

In the dorsolateral prefrontal cortex, Alzheimer’s disease is distinguished from both normal ageing and amyloid-positive, tau-negative pathology aging by (1) a progressive loss of neuropeptide and hormone-signalling transcripts, (2) an immune programme centred on interferon-γ responses, myeloid activation and classical complement that is already evident at the pathology-aging stage, and (3) an AD-specific induction of gliogenesis, oligodendrocyte differentiation and myelination genes together with the microglial receptor *P2RX7*. Fifty genes changed concordantly in both AD contrasts and form a high-confidence core signature. *APOE*- and *TREM2*-stratified analyses were uninformative owing to confounding and low power. These findings generate specific, testable hypotheses about stage-specific mechanisms in AD, and should be validated in larger, covariate-annotated, cell-type-resolved datasets before translation. Future work should therefore prioritize larger, balanced cohorts, covariate-aware models, and single-nucleus or spatial validation to assign these changes to specific cell types and disease stages.

## Declarations

### Data availability

The raw sequencing data are publicly available in the NCBI Sequence Read Archive under BioProject PRJNA1017546 (study SRP460672; samples GSM7782916-GSM7782930). Differential expression and enrichment tables generated in this study are available from the corresponding author on reasonable request.

### Ethics statement

This study is a secondary analysis of de-identified, publicly released human post-mortem data; no new ethical approval was required.

## Acknowledgements

We thank the brain donors and their families, and the investigators who generated and shared the underlying dataset.

