## Supplementary Figures 1-8 for "Prefrontal cortex transcriptomes distinguish Alzheimer’s disease from asymptomatic amyloid pathology, neuropeptide loss, immune-complement activation and glial remodelling"

Supplementary Figures S1–S8 present the remaining differential expression and enrichment analyses referred to in the Results.

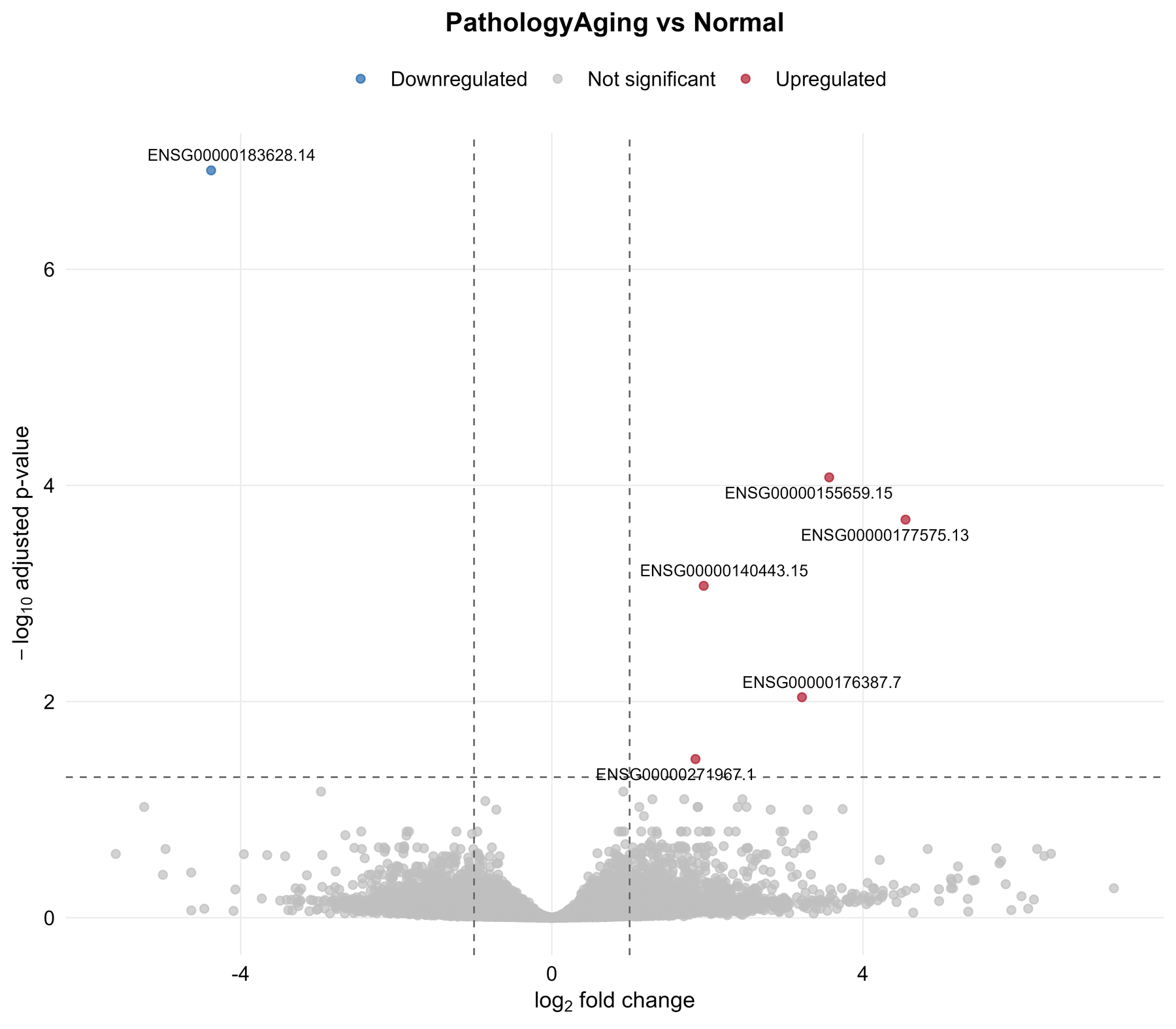

**Figure S1. Volcano plot of differential expression, pathology aging versus Normal.** Axes, thresholds and colours as in Figure 2. Only six genes met adjusted *P* < 0.05 and |log₂FC| > 1 (five up, one down; Table 5). Positive log₂FC denotes higher expression in PA.

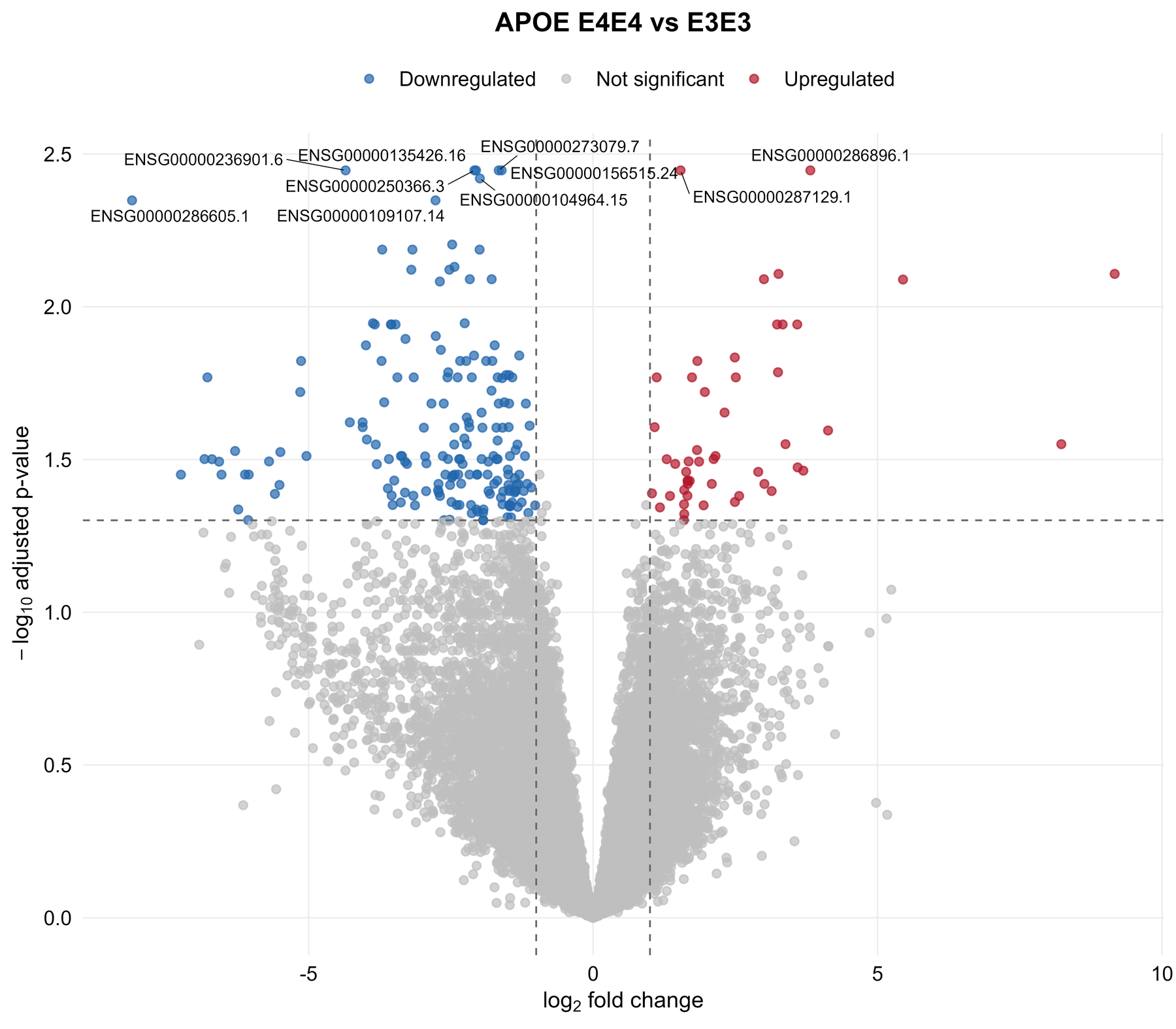

**Figure S2. Volcano plot of *APOE* E4/E4 versus E3/E3 donors (exploratory).** Positive log₂FC denotes higher expression in E4/E4 (*n* = 2) relative to E3/E3 (*n* = 5). The 235 DEGs (50 up, 185 down) are confounded with diagnosis because both E4/E4 donors are AD cases (Table 9).

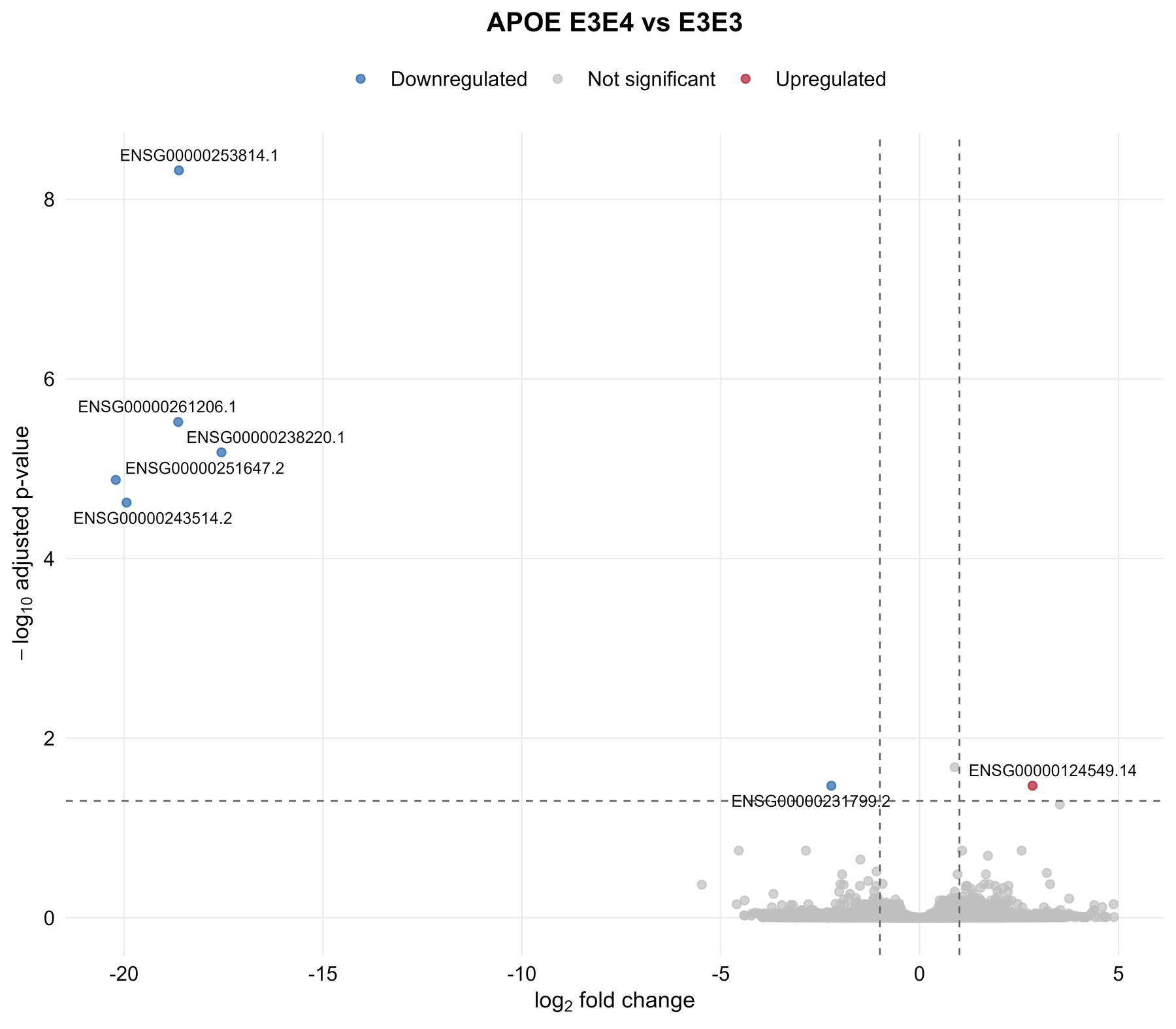

**Figure S3. Volcano plot of *APOE* E3/E4 versus E3/E3 donors (exploratory).** Positive log₂FC denotes higher expression in E3/E4 (*n* = 8) relative to E3/E3 (*n* = 5). Seven DEGs were detected, all non-coding or pseudogenes, most with mean normalised counts near one (Table 9).

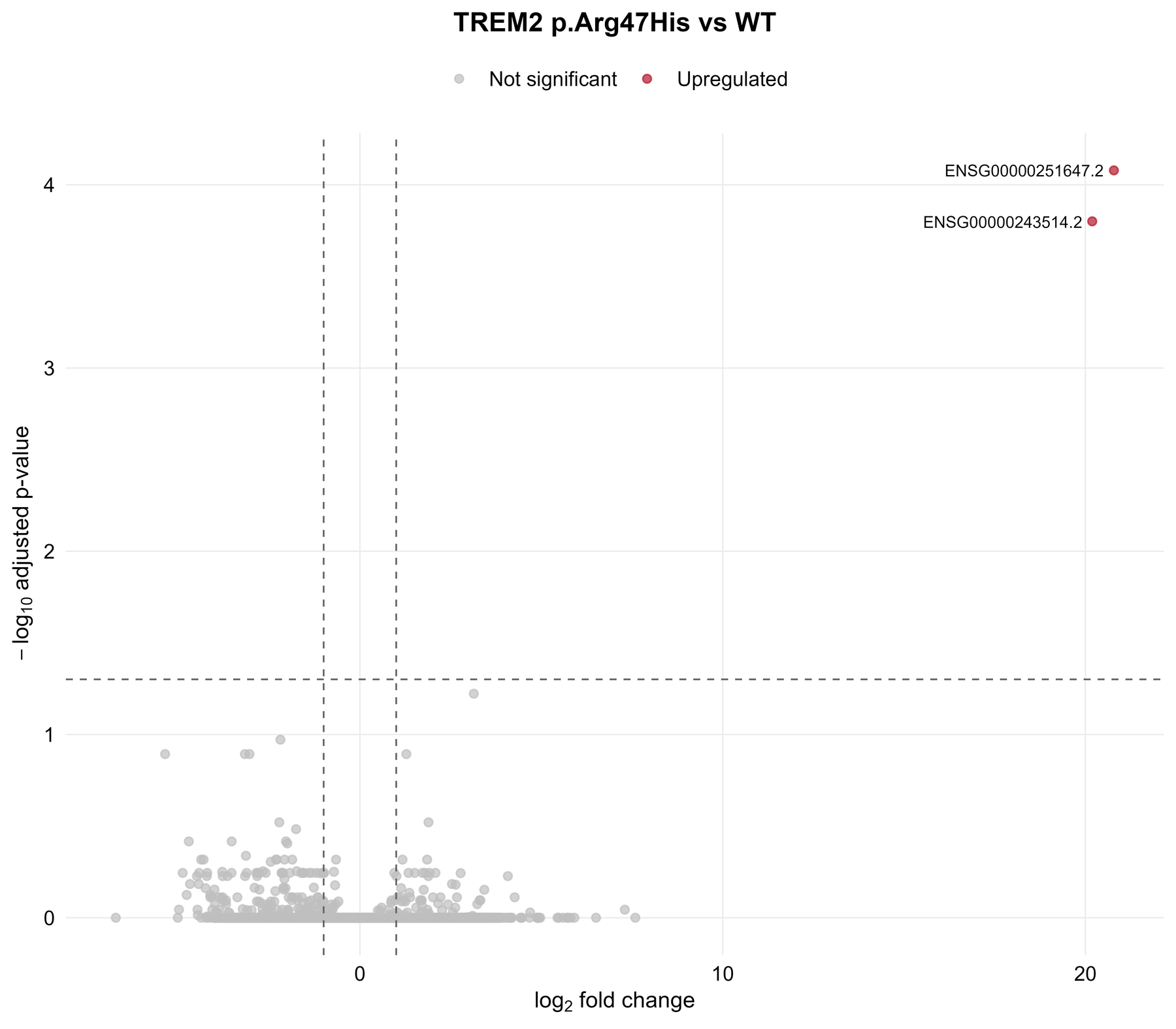

**Figure S4. Volcano plot of *TREM2* p.Arg47His carriers versus wild type (exploratory).** Positive log₂FC denotes higher expression in R47H carriers (*n* = 4) relative to wild type (*n* = 11). Two pseudogene DEGs were detected with mean normalised counts of about one (Table 9).

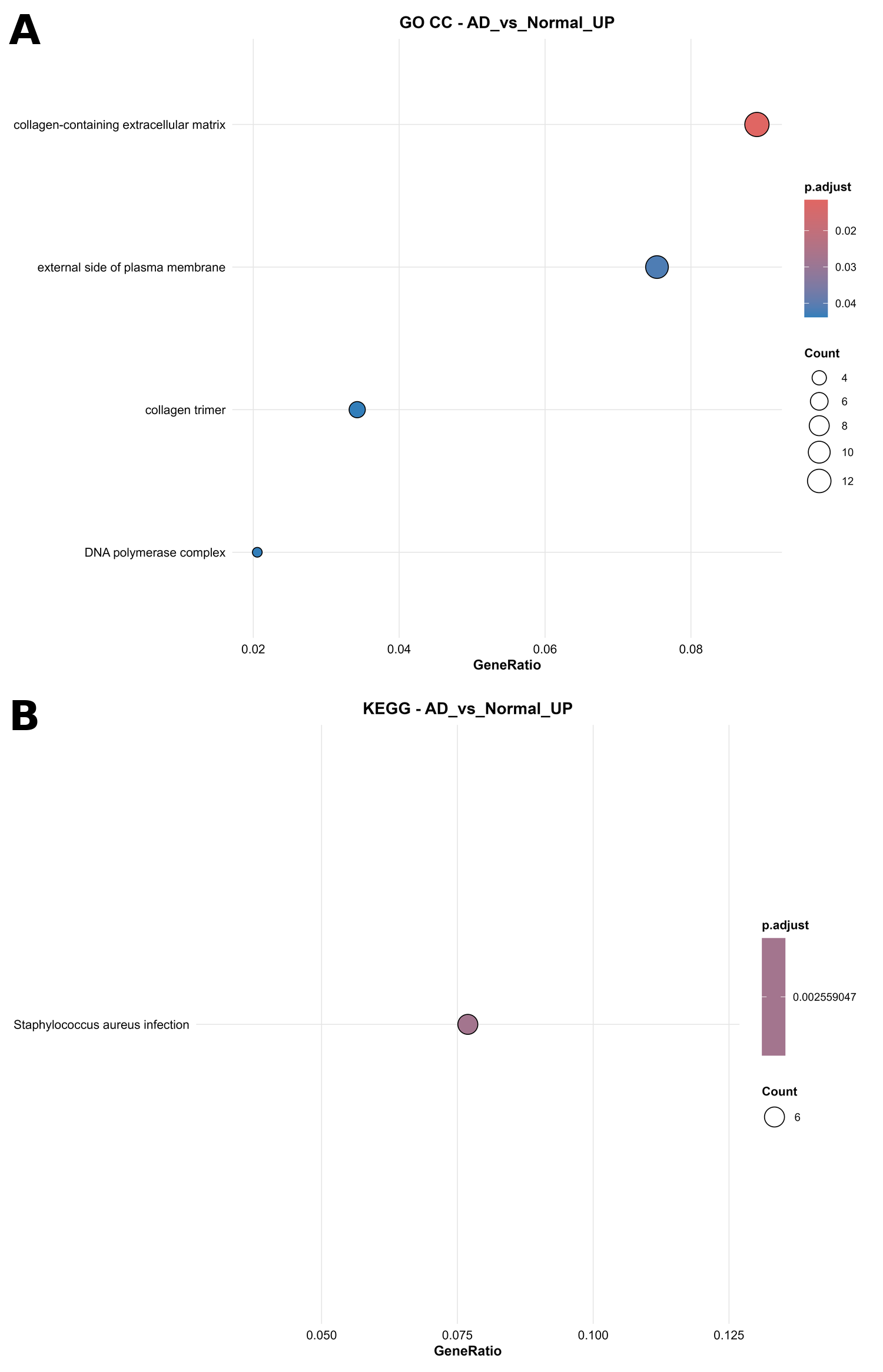

**Figure S5. Additional enrichment results for genes up-regulated in Alzheimer’s disease versus Normal.** (A) GO cellular component (collagen-containing extracellular matrix, external side of plasma membrane, DNA polymerase complex, collagen trimer); (B) KEGG pathways (*Staphylococcus aureus* infection). Plot encoding as in Figure 4.

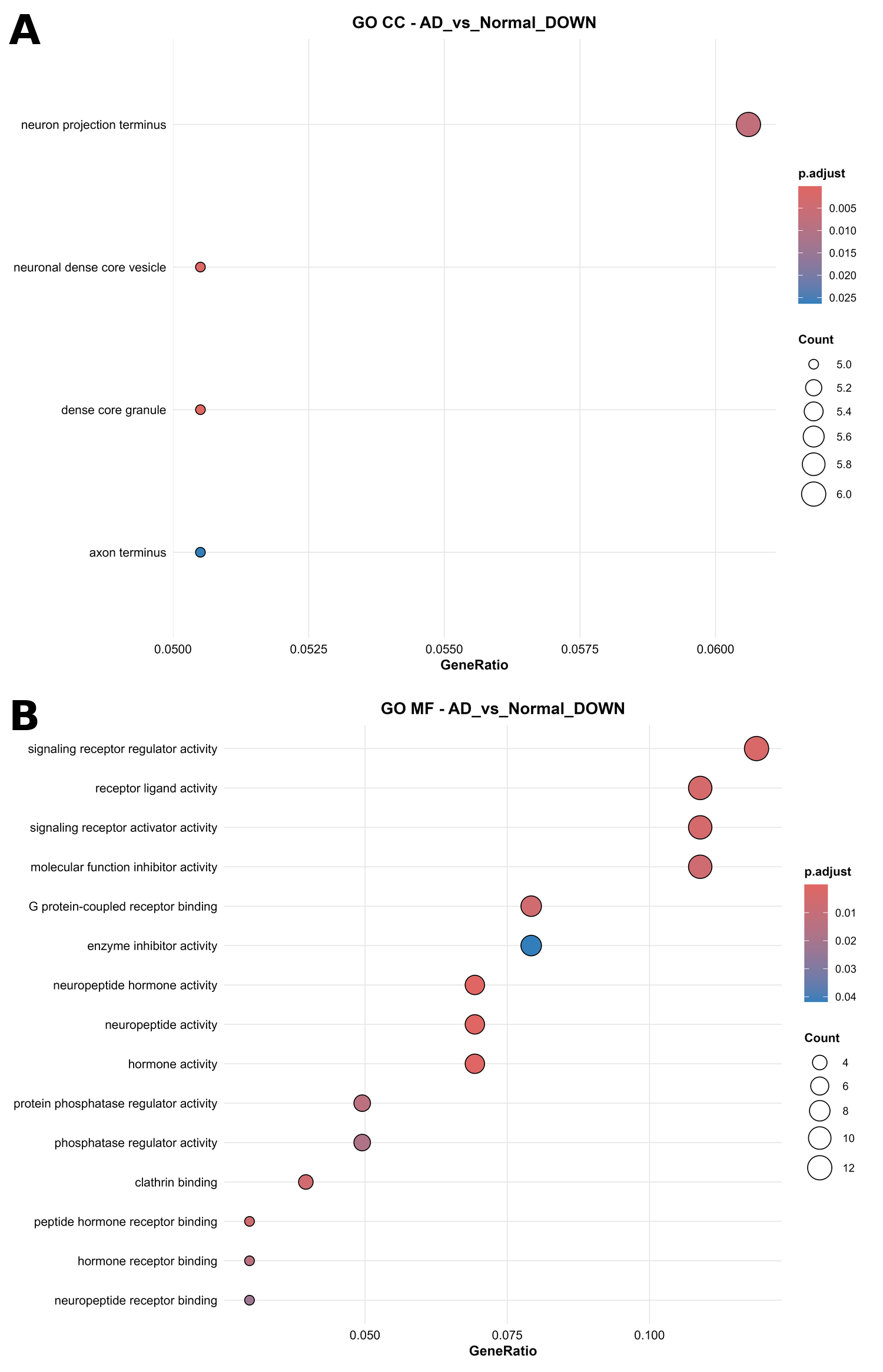

**Figure S6. Additional enrichment results for genes down-regulated in Alzheimer’s disease versus Normal.** (A) GO cellular component (neuronal dense core vesicle, dense core granule, neuron projection terminus, axon terminus); (B) GO molecular function (neuropeptide hormone activity, neuropeptide activity, hormone activity, receptor-ligand activity). Plot encoding as in Figure 4.

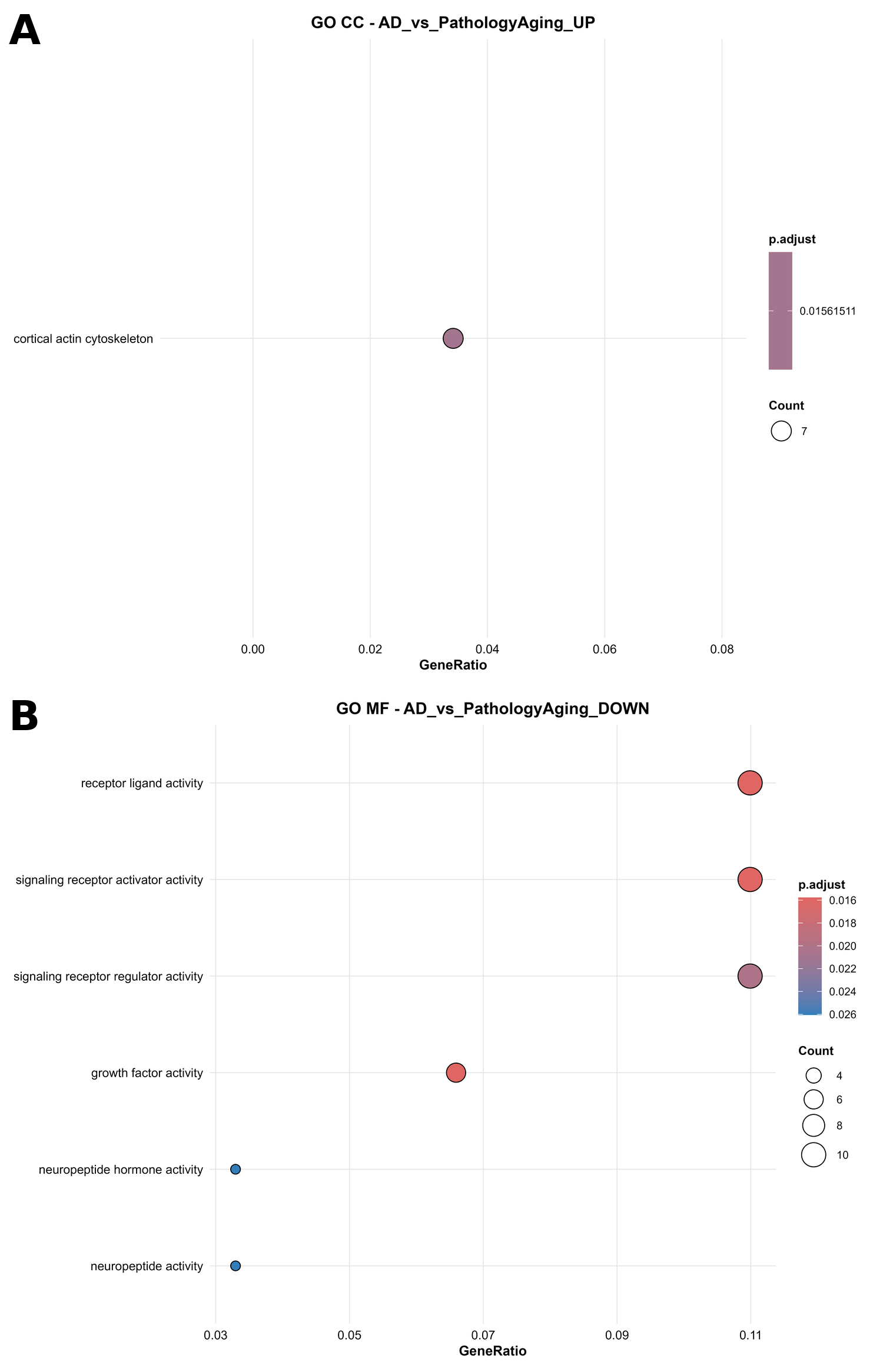

**Figure S7. Additional enrichment results for Alzheimer’s disease versus pathology aging.** (A) GO cellular component for up-regulated genes (cortical actin cytoskeleton); (B) GO molecular function for down-regulated genes (growth factor activity, receptor-ligand activity, neuropeptide hormone activity). Plot encoding as in Figure 4.

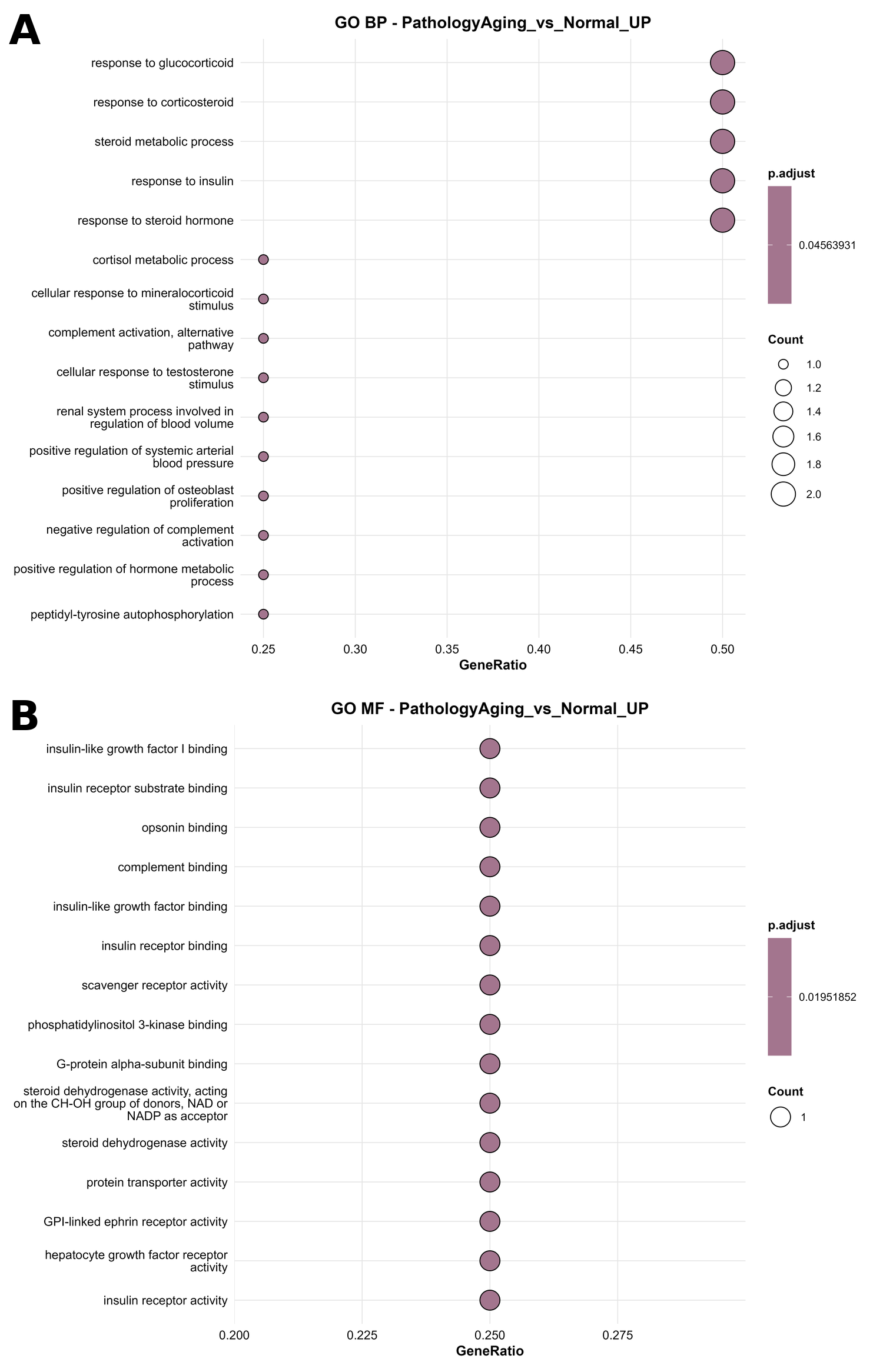

**Figure S8. Enrichment results for genes up-regulated in pathology aging versus Normal.** (A) GO biological process; (B) GO molecular function. Terms are driven by one or two of the five up-regulated genes (gene ratios of 1/4 or 2/4) and should be interpreted as descriptive (Table 8). Plot encoding as in Figure 4.
